# Multimodal temporal mapping of macrophage transcriptome remodeling during *Salmonella* infection

**DOI:** 10.64898/2026.07.04.736507

**Authors:** Christophe Toussaint, Peter W S Hill, Bastian Klodewig, Matthew Eldridge, Fabian J Theis, Sophie Helaine, Antoine-Emmanuel Saliba

## Abstract

Macrophages are equipped to eliminate invading pathogens, yet several intracellular bacteria exploit them as replicative niches. *Salmonella enterica* serovar Typhimurium subverts host immunity by injecting effector proteins that remodel macrophage functions. While macrophages typically induce a pro-inflammatory program upon bacterial invasion, *Salmonella* can redirect them toward an anti-inflammatory and replication-permissive state via manipulation of the NF-κB and STAT3 host transcription factors. How the integration of the effects on these two transcription factors and potentially others underpins this reprogramming remains poorly charted. Here, we use a multipronged approach combining a bacterial reporter, temporal single-cell RNA-seq with RNA metabolic labeling, transcription factor (TF) footprinting, and single-cell CRISPR perturbations to dissect macrophage polarization dynamics during early infection. We catch the bifurcation during infection, where a subset of macrophages transition toward the anti-inflammatory phenotype. This shift involves the activation of *Salmonella* pathogenicity island 2 (SPI2) leading to both the dampening of the initial NF-κB–driven inflammatory program and the induction of specific transcriptional modules beyond NF-κB and STAT3, with possible contributions from AP-1 and Maf family members. Together, our study uncovers host decision points in macrophage polarization circuitry and reveals a vulnerability exploited by *Salmonella* to modulate host immunity.

**HIGHLIGHTS:**

- Temporal scRNA-seq defines the emergence of inflammatory and anti-inflammatory macrophage states concomitant with *Salmonella* Typhimurium SPI2 induction.
- A critical decision point for infection outcome is reached by 6 hours post-infection.
- Macrophage reprogramming is associated with widespread chromatin remodeling and dynamic transcription factor binding.
- Single-cell perturb-seq experiment uncovers *Cybb* and *Foxo1* as modulators of infection outcome in Hoxb8 macrophages.

## INTRODUCTION

A large fraction of human intracellular bacterial pathogens use phagocytic cells, particularly macrophages, as preferred hosts for replication and persistence^1–4^. Therefore, elucidating the host factors and pathways that enable bacteria to evade antimicrobial macrophage functions^3^, modulate macrophage immune responses^1^, and create a metabolically permissive host^5^ is key to developing host-directed therapies to intercept the spread of infection^6,7^.

Macrophages display a wide repertoire of polarization states depending on the activation signals they receive^8^. Whereas macrophages typically assume a pro-inflammatory polarization state upon sensing bacterial invasion, *in vivo* macrophage states that sustain the intracellular survival of pathogens show features of alternatively activated and foamy macrophages, broadly defined as states characterized by nutrient-rich metabolic profiles and low inflammatory signaling. Prototypical examples include *Brucella abortus* and *Salmonella* Typhimurium (*S*Tm), which rely on glucose availability in macrophages to grow. Mice lacking the glucose metabolic regulators peroxisome proliferator-activated receptor (PPAR-γ) and -δ show decreased bacterial load^9,10^. These findings are supported *in vitro*, as interleukin-4 (IL-4)-stimulated bone-marrow-derived macrophages (BMDMs) that adopt an alternatively activated state can dramatically boost bacterial proliferation^9,10^. Yet it was only more recently revealed that bacteria do not only take advantage of a permissive environment but also actively shape permissiveness. Thus, STm actively reprograms macrophages via bacterial effectors of the pathogenicity island 2 (SPI2)-encoded type 3 secretion system (T3SS)^11^. Several effectors converge on reducing NF-KB activation and thereby dampen the pro-inflammatory response^12,13^ while SteE mediates the expression of STAT3-dependent genes^14^, eliciting an anti-inflammatory programme reminiscent of an M2-like polarization state. The reprogramming by SteE has recently been dissected and occurs at a genome-wide level, involving the activation of >1,000 genes and >1,000 chromatin regions with altered accessibility^14^. Because STm-dependent reprogramming of macrophages results from the integration of several effectors acting concomitantly, we sought to unravel the host transcriptional cascade that determines macrophage fate.

Macrophage transcriptional changes are asynchronous and heterogeneous across cells due to bacterial heterogeneity in SPI2 induction and bacterial growth. In addition, traditional RNA-seq, which relies on total RNA measurements, has poor temporal resolution^15^. We thus elected to resolve the gene network of inflammatory changes during STm infection using single-cell nascent RNA measurements^15,16^. Single-cell RNA-seq (scRNA-seq) with metabolic RNA labelling enables time-resolved monitoring of transcriptional responses for thousands of genes^17^. In particular, scSLAM-seq^18,19^, based on exposing cells to the nucleoside analogue 4-thiouridine (4sU), allows segregation of total RNA reads into old and new RNA molecules on the basis of T-to-C mismatches. RNA metabolic labeling thereby offers the opportunity to distinguish direct from indirect transcriptional effects of a perturbation, allowing more precise dynamic network reconstruction^20^.

Here, we combined temporal scSLAM-seq with a bacterial transcriptional reporter of SPI2 activation during STm macrophage infection. This allowed us to define SPI-2-dependent host gene modules involved in dampening the initial NF-κB–driven inflammatory program and activating the anti-inflammatory state. We then used transcription factor (TF) footprinting and single-cell CRISPR perturbations to resolve macrophage reprogramming dynamics and nominate candidate host regulators of the reprogramming trajectory beyond NF-KB and STAT3, with possible contributions from AP-1 and Maf family members. Together, these approaches reconstruct the transcriptional cascade that determines macrophage fate and begin to resolve when, and in which cells, *Salmonella* is able to impose it.

## RESULTS

### scSLAM-seq captures the macrophage response dynamics tied to *Salmonella* activity

The transcriptional dynamics of the macrophage response to infection and subsequent bacterial-driven reprogramming occur during the first ten hours of STm infection. We therefore designed a temporal single-cell RNA-seq workflow integrating a bacterial SPI2 readout, RNA metabolic labelling, single-cell fluorescence-based sorting on plates, and single-cell RNA-seq, sampling every two hours over the first ten hours, with additional timepoints at 18 and 21 hours of infection as controls comparable to datasets from previous studies^11,21^ (**Figure 1A**). Building on our previously established infection model^21^, we infected murine bone-marrow derived macrophages (BMDMs) with a *S*Tm strain carrying a plasmid reporting on the growth history of internalized bacteria through the expression of two fluorescent proteins. A constitutively expressed mCherry allowed the discrimination of infected macrophages and served as a proxy for intramacrophage bacterial load over time. An unstable variant of GFP under the control of the *ssaG* promoter functioned as a reporter of SPI2 induction. Flow cytometry enabled the sorting of infected macrophages into three groups: cells harbouring SPI2-inactive bacteria, SPI2-active non-growing bacteria, and SPI2-active growing bacteria (**Figure 1B**). In line with previous reports, SPI2 induction started between 4 and 6 h of infection, and bacterial growth followed between 6 and 8 h (**Figures S1A and S1B**).

**Figure 1:**
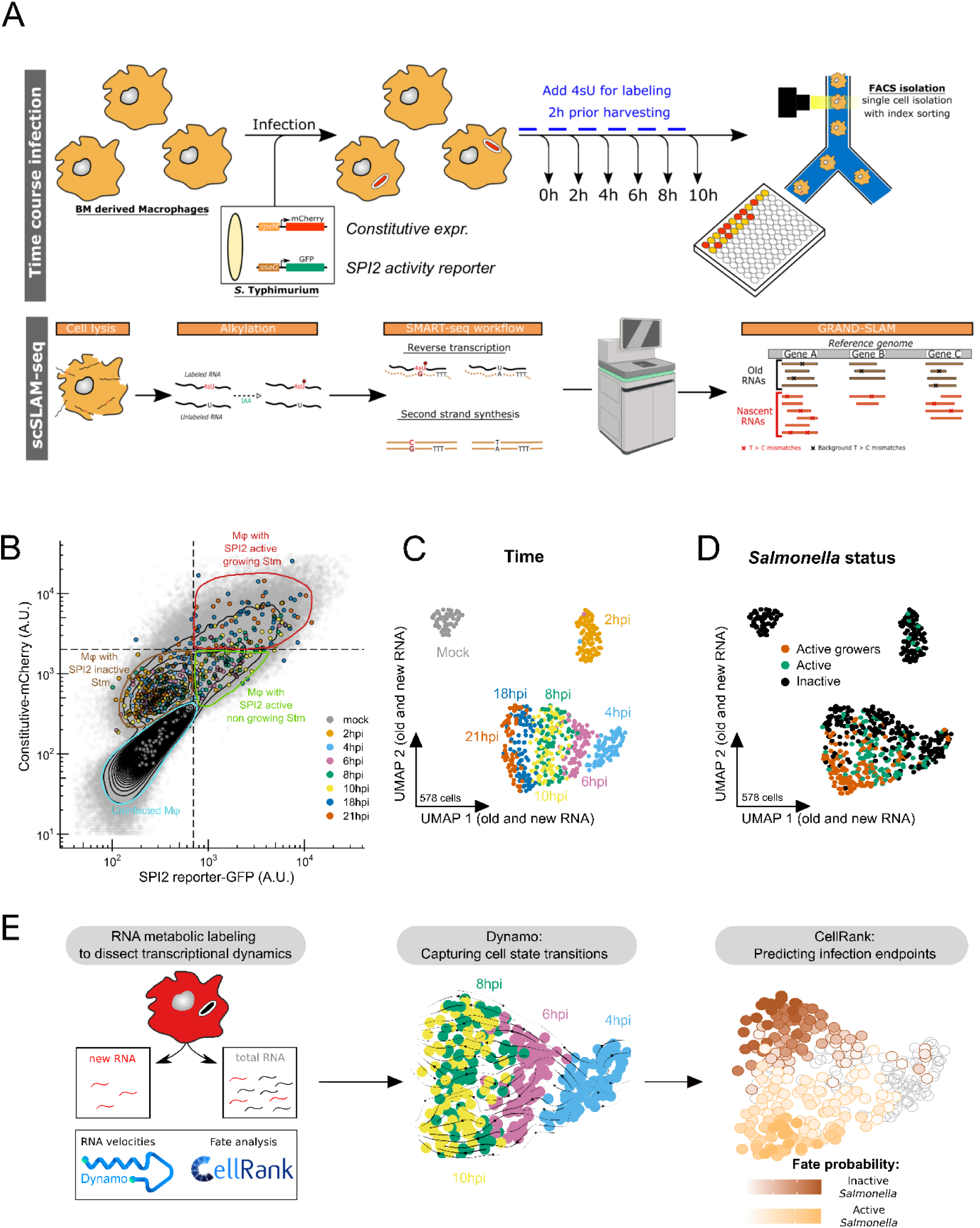
Connected macrophage fate trajectories to Salmonella SPI-2 activity through scSLAM-seq. **(A)** Experimental (upper panel) and analytical (bottom panel) workflow depicting the time-course and multimodal scSLAM-seq analysis of infected bone marrow derived macrophages with Salmonella carrying a SPI2-reporter plasmid. **(B)** Aggregated scatter plot of intracellular bacterial SPI2 activity in non-infected (mock) and infected macrophages analysed and sorted by FACS across the complete time-course of the experiment time points of infection analysed. Grey background data points and contour lines correspond to the entire cell population analysed superimposed with timepoint-colored dots indicating all the individual sorted macrophages for downstream analysis color-coded by time-point further processed for scSLAM-seq. The scatter plot has been segmented in four quadrants based on bacterial intracellular activity. **(C & D)** UMAP (Uniform Manifold Approximation and Projection) visualization of 578 single-macrophage transcriptomes obtained by scSLAM-seq. UMAP embeddings result from the joint analysis of new and old RNA levels. Single macrophages are color-coded by timepoint of collection (C) and Salmonella SPI2 activity (D) as inferred from mCherry and GFP fluorescence levels recorded by flow cytometry. **(E)** Bioinformatic workflow to leverage RNA metabolic labeling information for transcriptome dynamics analysis and cell fate prediction (Left). Focusing on 4 hpi to 8 hpi single-cell embedding at the onset of SPI2 activity (panel C/D), a velocity analysis (middle) and fate probability has been carried out (right).

As macrophages undergo complex remodelling, RNA metabolic labelling increased our power to detect genes undergoing transcription variations on short time scales. 4sU-labeled RNA was alkylated with iodoacetamide (IAA), causing characteristic T-to-C mutations during reverse transcription. These mutations enabled sequencing-based distinction between newly synthesized and pre-existing RNA, allowing estimation of gene-specific new-to-total RNA ratios (NTR)^22^ (see **Methods**). We sequenced a total of 836 single-cell transcriptomes, including 652 cells treated with 4sU. After quality control, we kept a total of 765 single cells meeting high quality metrics with respect to the number of detected genes (median of 4,662 genes per cell) and percentages of exonic, mitochondrial, and ERCC spike-in reads (**Figure S1C**). We finally retained 578 cells with effective 4sU labelling for analysis relying on new RNA levels **(see Table S1)**. Altogether, our approach therefore recorded for each macrophage five layers of information: time, old RNA measurements, new RNA measurements, SPI2 expression, and growth status.

To explore the scRNA-seq data, we decomposed new RNA and old RNA count matrices before making joint use of both for dimensional reduction, visualization and identification of macrophage transcriptomic states (**Figures 1C, 1D; Methods**). A first inspection of the UMAP showed that macrophages segregated by infection timepoint, with the changes becoming less pronounced from 8 hpi onwards (**Figure 1C**). This visualization also revealed an increase in the heterogeneity of transcriptomic states displayed by BMDMs starting at 6 hpi (**Figure 1C**).

Notably, this rise in heterogeneity coincided with the activation of SPI2 expression between 4 and 6 hpi in a subpopulation of internalized *S*Tm and remained a discriminating marker at later timepoints (**Figure 1D, Figures S1A**). We confirmed that 4sU exposure did not interfere with mock or infected macrophages over the time course, as 4sU-treated and untreated cells were not segregated (**Figure S1D**). T-to-C and A-to-G mismatch frequencies were markedly increased in all cells exposed to 4sU compared to untreated controls (**Figure S1F**). Additionally, no batch effect between biological replicates could be detected (**Figure S1E**). Global percentages of new RNA per single-cell varied across timepoints (median of 29.6%), with a peak in the proportion of new RNA at 4 hpi (**Figure S1G**).

To further leverage the temporal information captured by RNA metabolic labelling, we next set up an analysis framework based on RNA velocities to reconstruct cell state transitions and compute cell fate probabilities^23,24^ (**Figure 1E**). From the velocities, we could identify, starting at 4 hpi, two divergent trajectories that matched bacterial SPI2 activity (**Figure 1D**). Altogether, having used our multimodal data to reveal the transcriptional dynamics of macrophage remodelling, we next set out to characterise the different macrophage states arising over time.

### Macrophage polarization is not fixed by the early response

We then sought to characterize the diversity of macrophage states emerging during *S*Tm infection. Using unsupervised clustering with the Leiden algorithm, we identified 11 distinct macrophage states (**Figure 2A**). These states largely recovered the infection timepoints up to 4 hpi, suggesting a stereotypical early transcriptional response. In contrast, from 6 hpi onwards, multiple macrophage states were observed within each timepoint, coinciding with increasing transcriptomic asynchrony and aligning with differences in *S*Tm SPI2 activity (**Figures 1C, 1D and 2A**).

**Figure 2:**
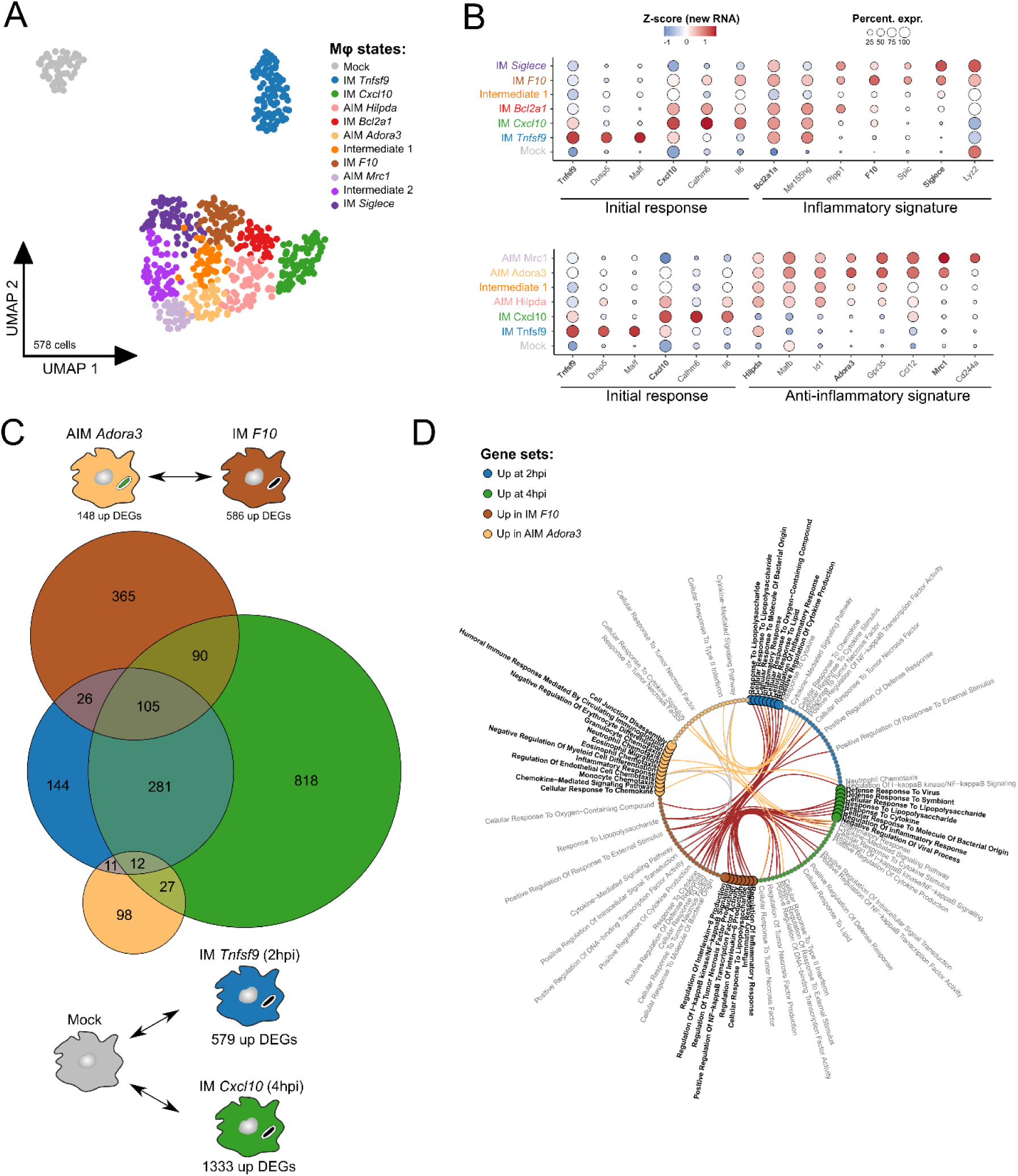
Dynamics of transcriptional response of macrophages. **(A)** UMAP visualization of 578 single-macrophage transcriptomes obtained by scSLAM-seq as in Figures 1C and **1D** and colored by cell state as identified by unsupervised clustering with the Leiden algorithm. **(B)** Dot plot of scaled, log-normalized expression of marker selected representative genes from the different macrophage states identified (A). Top and bottom panels correspond to the inflammatory and anti-inflammatory macrophage trajectory, respectively. Dot size indicates the percentage of cells per cell state with detectable mRNA expression, and color shows Z-scores of log-normalized new RNA readcounts. **(C)** Analysis of differentially expressed genes overlap between different macrophage states represented by a Venn plot of differentially expressed genes (FDR < 0.05, Wilcoxon Rank Sum test with Benjamini-Hochberg correction) differentiating the early response at 2/4hpi and the inflammatory/anti-inflammatory signatures. Circle size is proportional to the total number of DEG within each group. Indicated numbers correspond to the number of specific or shared DEGs between groups. Comparisons between macrophage states performed to identify each DEG group are represented (top and bottom). **(D)** Edge bundling graph of pathway enrichment analysis results for each DEG group presented in (C). Edges depict enriched pathways either from inflammatory or anti-inflammatory signatures that are common with the early response. Top 8 enriched pathways per DEG group are indicated in black. Other significant pathways shared between DEG groups are indicated in grey (FDR < 0.05, Fisher exact test with Benjamini-Hochberg correction).

All macrophage states were annotated based on differential gene expression (DEG) analysis between states, across and within timepoints (**Figure 2A and 2B**). First, they were broadly classified as inflammatory (IM) or anti-inflammatory (AIM) based on their overall transcriptional signatures (**Figures S2A**). Then, representative genes showing state-specific or peak expression were used to further refine each cluster annotation (**Figure 2A**). We also identified two intermediate (INT) states exhibiting mixed transcriptional features of both inflammatory and anti-inflammatory macrophages (**Figures 2A and 2B**).

We then focused on the inflammatory and anti-inflammatory populations observed at 8 and 10 hpi (IM *F10* and AIM *Adora3*). These states were homogeneous with respect to *S*Tm activity (**Figure 1D**) and comprised cells originating from different timepoints, suggesting that they represent relatively stable transcriptional programs. We therefore performed DEG analysis to define their molecular signatures. Overall, DEG testing revealed a gradual and sequential activation of gene programs leading to the emergence of these states at 8/10 hpi and persisting up to later timepoints (18-21 hpi) (**Figures 2B and S2A**). IM *F10* and AIM *Adora3* displayed increased expression of canonical markers previously identified in late BMDM infection of M1 and M2 polarization^11,21^, respectively, including *Tnf* and *Il1b* for M1, and *Mrc1* (*Cd206*), *Arg1*, and *Il4ra* for M2 (**Figures S2B and S2C**). These observations support the designation of these states as inflammatory and anti-inflammatory while also revealing a more nuanced picture of a spectrum rather than a binary transcriptional state.

We then examined the early transcriptional response occurring within the first hours of infection (up to 4 hpi). We observed different patterns of gene expression, including transiently induced genes restricted to specific timepoints (e.g. *Dusp5* at 2 hpi and *Calhm6* at 4 hpi), as well as genes associated with NF-κB signalling (e.g. *Bcl2a1* family members and *Mir155hg*) that were strongly induced early and remained elevated in inflammatory macrophages (**Figure S2A**).

We then assessed to what extent the inflammatory and anti-inflammatory signatures at 8/10 hpi overlapped with genes induced during the early response (**Figure 2C**). A fraction of DEGs from both inflammatory and anti-inflammatory signatures were part of the initial macrophage response at 2 and 4 hpi, consistent with a model in which an initial inflammatory response also primes a potential negative feedback loop. Notably, module scores derived from the full set of early response genes did not distinguish between inflammatory and anti-inflammatory macrophages at 8–10 hpi (**Figure S2B**), highlighting the importance of more specific gene programs in defining these states.

To place these transcriptional patterns in a functional context, pathway enrichment analysis revealed that the early response was dominated by inflammatory processes, with Gene Ontology terms related to pathogen response, cytokine production, and NF-κB signaling (**Figure 2D**). While the IM *F10* signature largely overlapped with this profile, the AIM *Adora3* signature instead highlighted processes linked to immune cell chemotaxis and migration, pointing to a distinct transcriptional program.

Together, these results show that we captured the changes accompanying STm SPI-2-dependent reprogramming of macrophages from the early pro-inflammatory response into an anti-inflammatory state.

### RNA metabolic labelling uncovers transcriptional dynamics that lead to the establishment of the inflammatory and anti-inflammatory states

Having established that IM *F10* and AIM *Adora3* states represent relatively stable polarization signatures, we sought to dissect the sequence of transcriptomic events leading to those states after the early response of macrophages to *S*Tm.

First, we identified four main gene modules by testing for differentially expressed genes between IM *F10* and AIM *Adora3* states. Macrophages from the intermediate state were also included in the analysis to capture more subtle transcriptomic differences (see **Methods**). Genes within each module could be further grouped based on their pattern of induction across the infection: genes induced early in the infection, up to 4 hpi (i.e. significantly upregulated in IM *Tnfsf9* or IM *Cxcl10* states compared to uninfected macrophages); and genes with delayed induction beyond the 6 hpi timepoint (**Figure 3A**, heatmap). Irrespective of induction pattern, each module correlated with SPI2 expression (**Figures S3A and S3B**).

**Figure 3:**
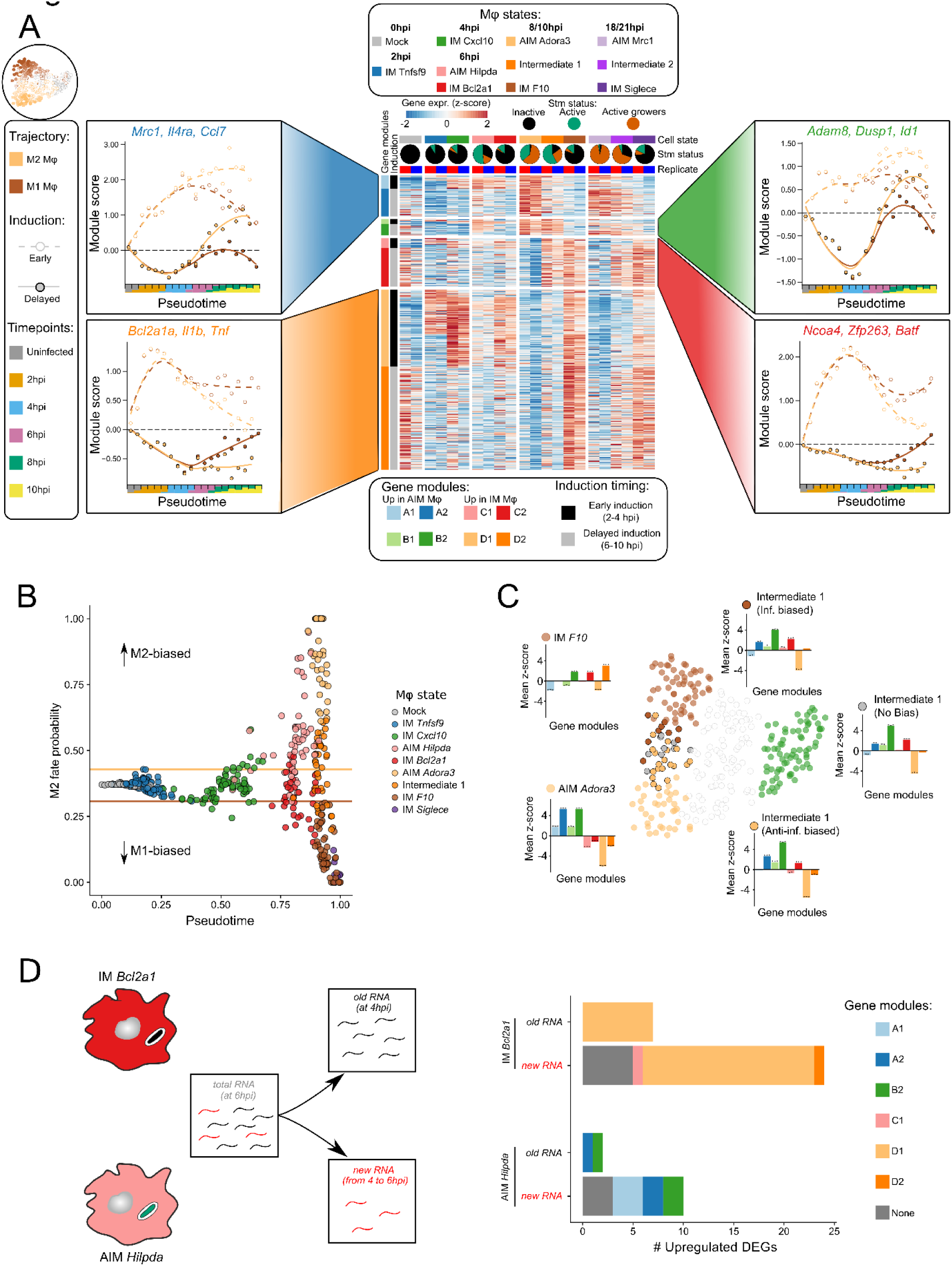
RNA metabolic labeling uncovers transcriptional dynamics leading to the establishment of the Inflammatory and Anti-inflammatory signatures. **(A)** (Middle) Heatmap displaying z-scores of log-normalized new RNA readcounts of differentially expressed (DE, FDR < 0.05) genes between Inflammatory and Anti-inflammatory macrophage states at 8/10hpi. Single macrophage transcriptomes have been pooled per biological replicate (n = 2) for each macrophage state. Pie charts represent the proportion of Salmonella status within a given macrophage state across both replicates as defined in Figure 1B. DE genes were classified into a total of 8 modules based on their specific expression to Inflammatory or Anti-inflammatory cell states or shared expression with partially reprogrammed macrophages (4 main modules) and timing of DE gene induction following infection (early or delayed). Statistical significance of differential expression for each gene per macrophage state is shown in Table S2. (Left and right) Module score trend plots along Inflammatory and Anti-inflammatory trajectory pseudotime. Module scores were calculated based on the log-normalized new RNA readcounts and each panel corresponds to one of the main gene modules including the indicated representative genes. Datapoints correspond to the module score weighted mean based on CellRank fate probabilities for each bin of the trajectory pseudotime (1 bin = 22 or 23 cells). The proportion of collection timepoints per bin is indicated on the x-axis. Dashed and solid color lines correspond to the timing of induction of DE genes within the module (respectively, early and delayed). Dashed black lines represent the baseline module scores computed in mock macrophages. CellRank fate probabilities and pseudotime values for each single macrophage are reported in Table S1. **(B)** Fate probability plot depicting the endpoint likelihood of each single macrophage. Each cell is ordered along the trajectory pseudotime computed with CellRank (x-axis) while y-axis correspond to cell fate probabilities towards Anti-inflammatory and Inflammatory endpoints as computed by CellRank (left and right y-axis respectively). Solid colored horizontal lines correspond to maximum and minimum fate probabilities for the IM Tnfsf9 macrophage state (2hpi). Data points are colored by macrophage state as indicated in Figure 2A. CellRank fate probabilities and pseudotime values for each single macrophage are reported in Table S1. **(C)** Gene module expression dynamics in Inflammatory, Anti-inflammatory and partially reprogrammed macrophages. Barplots depict mean z-score variations of gene modules identified in (A) for the indicated macrophage states relative to the M1 Cxcl10 macrophage state (4 hpi). Significant upregulations and downregulations are reported (*** FDR < 0.001, ** FDR < 0.01, * FDR < 0.05, One-sample t-test with Benjamini-Hochberg correction). UMAP embedding is zoomed on macrophages collected from 4 hpi to 10 hpi (300 cells) and cells linked to barplots are colored by cell state as in Figure 2A. **(D)** Results of DE gene testing between 6 hpi macrophage states in either old or new RNA levels (FDR < 0.05, Wilcoxon Rank Sum test with Benjamini-Hochberg correction).

To gain deeper insights into expression dynamics and link gene module activities to cell fate, we devised an approach specifically leveraging new and old RNAs (**Figure 1E**). We used RNA velocities, capturing the directionality of transcriptional changes with Dynamo^23^ from which we inferred a cell state transition matrix with CellRank^24^. We then calculated for each macrophage its probability of progressing toward inflammatory or anti-inflammatory states, and derived a pseudotemporal ordering along these trajectories. Gene module activity was quantified using module scores derived from new RNA levels. To capture module dynamics along trajectories toward inflammatory and anti-inflammatory states, mean module scores were visualized across the binned CellRank pseudotime (**Figure 3A**, side panels; see **Methods**).

Early-induced gene modules were upregulated in all infected macrophages irrespective of their classification. From 4hpi onward, their dynamics diverged between trajectories, with reciprocal suppression of opposing programs. Inflammatory modules (C1 and D1) peaked at 4 hpi and persisted at lower levels along the inflammatory trajectory, whereas anti-inflammatory modules (A1 and B1) were maintained or reinforced along the anti-inflammatory trajectory (**Figure 3A**). Conversely, delayed gene modules were initially downregulated relative to uninfected macrophages and progressively increased from 6hpi onward. Along the inflammatory trajectory these modules were restored to baseline, whereas along the anti-inflammatory trajectory the delayed inflammatory modules (C2 and D2) remained suppressed and the delayed anti-inflammatory modules (A2 and B2) were induced beyond baseline levels.

Next, we examined the relationship between CellRank pseudotime and cell fate probabilities to determine when infected macrophages committed to either trajectory (**Figure 3B**). Consistent with gene module dynamics, commitment toward inflammatory or anti-inflammatory states became apparent from 6hpi onwards (IM *Bcl2a1* and AIM *Hilpda* states). In contrast to the more polarized IM *F10* and *AIM* Adora3 states, intermediate macrophages at 8–10 hpi spanned a continuum of fate biases.

Given that Intermediate macrophages displayed a mixed transcriptional signature of inflammatory and anti-inflammatory modules, as well as heterogeneous STm SPI2 activity (**Figure 3A**, heatmap), we hypothesized that this population represented a partial reprogramming state reflecting ongoing reconfiguration of transcriptional programs prior to commitment. We therefore investigated how gene module activity is associated with fate bias within this population. To this end, we assessed the differential induction or repression of gene modules relative to the end of the early macrophage response (i.e. the IM *Cxcl10* state) (**Figure 3C**).

Irrespective of their eventual fate, intermediate macrophages displayed a mixed transcriptional program, lacking key features of both polarized states. Specifically, they failed to induce module D2, which characterizes the IM *F10* state, and did not exhibit the reinforced expression of module A1 observed in AIM *Adora3* macrophages. Nevertheless, the analysis suggests that intermediate macrophages commit through distinct routes, either coordinated upregulation of A1 coupled with downregulation of C1 along the anti-inflammatory path, or upregulation of C1 along the inflammatory trajectory. Taken together, these observations indicate that intermediate macrophages represent transitional states marked by partial and unresolved gene module activation and repression, and point to the C1 module as a candidate host decision point.

Finally, we focused on gene signatures associated with the commitment of infected macrophages toward inflammatory or anti-inflammatory outcomes as early as 6 hpi (**Figure 3B**). To this end, we performed differential expression analysis on the dominant macrophage states at this timepoint, considering both new and old RNA levels (**Figure 3D**). Notably, we detected only a limited number of DEGs based on old RNA levels, suggesting that the transcriptional shift toward the IM *Bcl2a1* and AIM *Hilpda* states occurs predominantly within the 4–6 hpi window. We next mapped the identified DEGs to their respective gene modules and found that genes upregulated in the IM *Bcl2a1* state predominantly belong to the early response module (D1). In contrast, the AIM *Hilpda* signature reflects a more heterogeneous contribution, encompassing both genes associated with the initial inflammatory response (A1) and genes from later-induced modules (A2 and B2).

Taken together, leveraging RNA metabolic labeling allowed us to resolve the dynamic expression of gene modules underlying inflammatory and anti-inflammatory macrophage states. While inflammatory macrophages sustained the early inflammatory response, albeit with reduced intensity after 4 hpi, SPI2-dependent reprogramming instead led to marked dampening of inflammatory gene modules and concomitant induction of an anti-inflammatory program.

### Transcription factor network analysis reveals shared regulatory control of macrophage polarization

We reasoned that the identified gene modules could be viewed as molecular fingerprints of upstream TF activities. We therefore sought to link the differentially expressed genes to their putative TFs using the TF-target regulons reported in the CollecTri database^25^ (see **Methods**). We identified three groups of transcription factors (TFs) based on their selective or shared connectivity to genes belonging to inflammatory and anti-inflammatory modules (**Figure 4**). Several of these TFs were also differentially expressed at the RNA level, including *Mxd1*, *Bach1*, and *Mafb* in association with anti-inflammatory modules, and *Nfkb2*, *Batf*, and *Nfe2l2* in association with inflammatory modules. The analysis also nominated candidate TFs that, despite not being differentially expressed, may still contribute to shaping gene module expression. Notably, *Stat3* was linked to both inflammatory and anti-inflammatory gene signatures, consistent with STAT3 being activated during infection through both the SteE/GSK3 anti-inflammatory axis^26,27^ and SteE-independent TLR4-dependent signalling with subsequent IL-6/IL-10 feedback as part of the general inflammatory response^28,29^. Similarly, components of the NF-κB pathway, including *Nfkb1* and *Rela*, were not exclusively associated with inflammatory gene modules. This association with both inflammatory and anti-inflammatory gene modules likely reflects the broad regulatory roles of STAT3 and NF-κB, whose target genes span both programmes.

**Figure 4:**
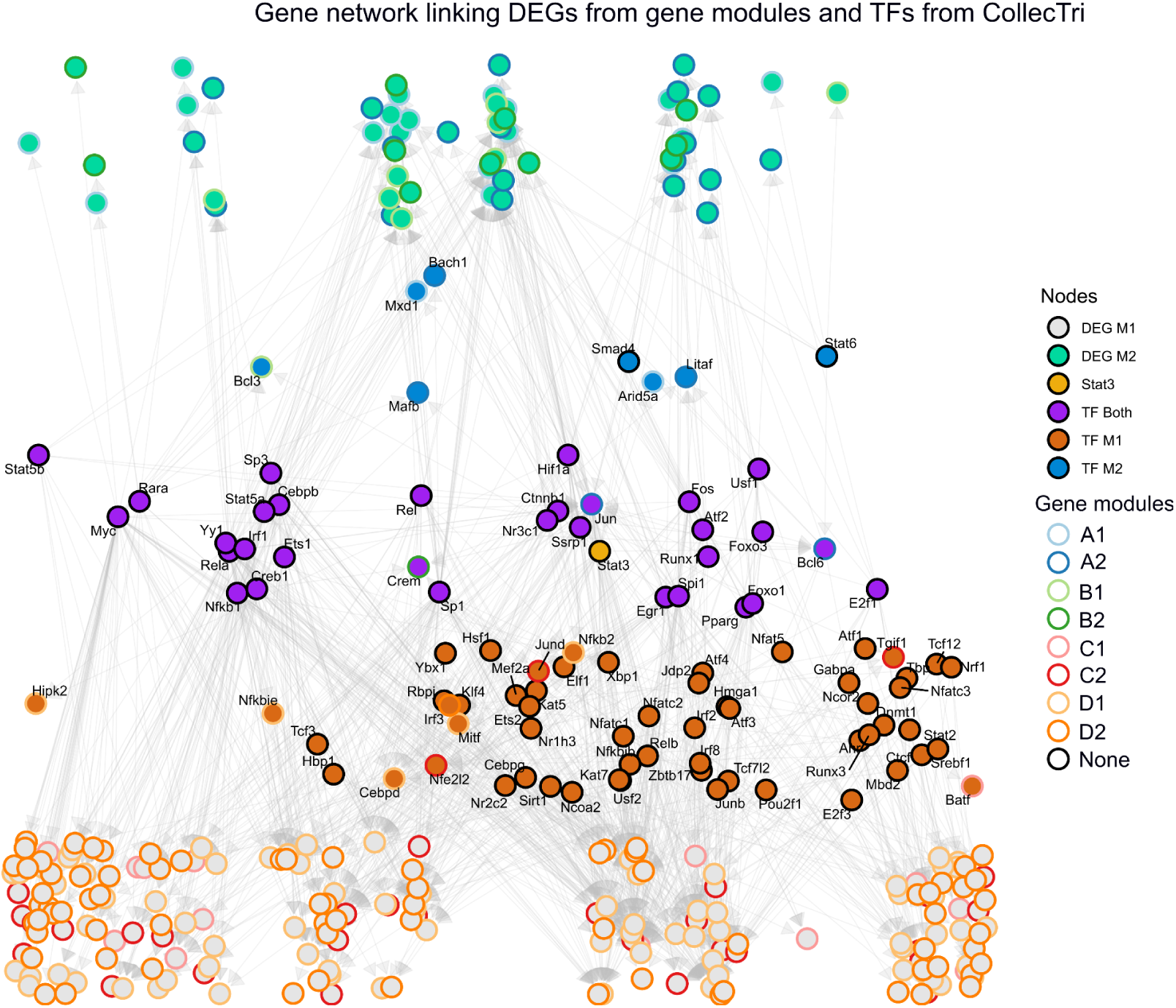
A map of upstream transcription factor regulating gene modules underlying macrophage polarization states. Transcriptional regulatory network linking TFs to DEGs in macrophages at 8–10 hpi. Networks were reconstructed from CollecTRI interactions by tracing upstream regulators of DEGs associated with inflammatory or anti-inflammatory modules. Nodes represent TFs or genes, and edges indicate regulatory relationships. Node colors denote TF class (inflammatory, anti-inflammatory, or shared) and DEG category, with Stat3 highlighted. Node borders indicate gene module assignment from Figure 3A. Only highly connected TFs and DEGs with upstream regulators are displayed.

Consequently, regulon-based analyses link these TFs to both modules but cannot determine which upstream signals or polarization states drive their activity in each context. Overall, our TF network analysis suggests that few TFs are purely specific to anti-inflammatory or inflammatory polarization and motivates direct functional perturbation to dissect their contributions.

### Transcription factor binding is redistributed between inflammatory and anti-inflammatory macrophage states

To investigate the involvement of the identified TFs in the establishment of inflammatory and anti-inflammatory macrophage polarization, we complemented our single-cell transcriptomic analysis with an ATAC-seq (Assay for Transposase Accessible Chromatin with sequencing) analysis. We performed bulk ATAC-seq and RNA-seq on matched samples of BMDMs infected for 2 or 18 hours with either wild-type (WT) STm or an isogenic Δ*ssaV* mutant unable to translocate SPI2 effectors into the host cell. Both strains carried the previously described SPI2 reporter plasmid, and infected macrophages were separated from uninfected cells by sorting on constitutive reporter expression. Because Δ*ssaV*-cannot use SPI2, the WT-infected BMDMs samples were additionally sorted for SPI2 activity. We compared transcriptomic signatures from the generated RNA-seq libraries to macrophage states identified in the scSLAMseq dataset (**Figure 5A**). Overall, we observed strong concordance between datasets: Δ*ssaV*-infected BMDMs closely resembled the pseudobulk of the IM *Siglece* state, whereas SPI2-active WT–infected BMDMs aligned with the AIM *Mrc1* state. We next analyzed the ATAC-seq profiles to assess differential chromatin accessibility at 18 hpi between Δ*ssaV*-infected BMDMs and those harboring SPI2-active WT STm. As expected, accessibility changes broadly correlated with gene expression (**Figure S4A**). Together, these observations support the use of the ATAC-seq dataset to explore activities of transcription factors associated with macrophage polarization during STm infection.

**Figure 5:**
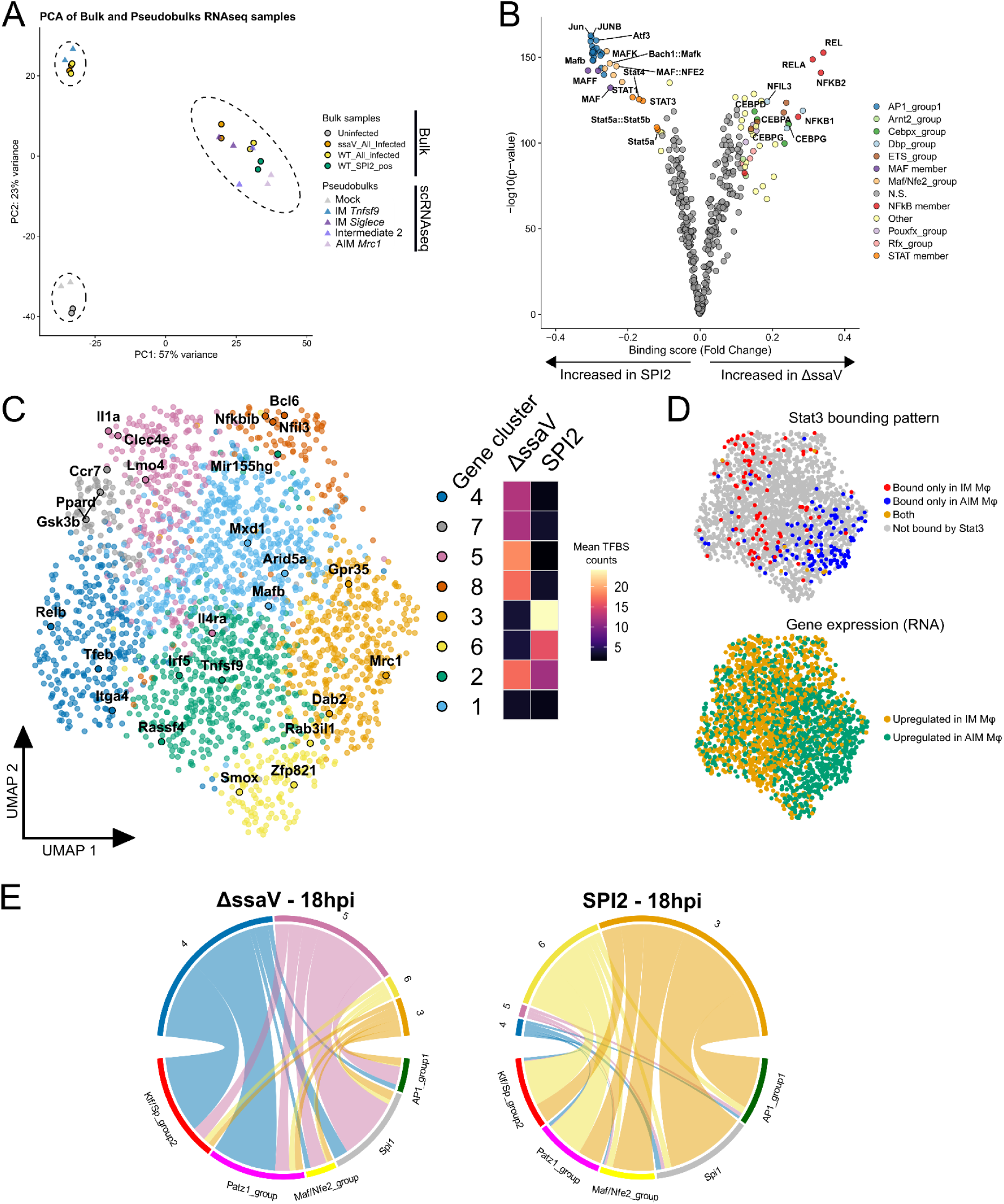
Transcription factor footprinting reveals wide TF repertoire shift. **(A)** Principal component analysis (PCA) performed on gene expression profiles from bulk RNA-seq samples (paired with ATAC-seq) and pseudobulks generated from the scSLAMseq dataset. Each point corresponds to one sample from one biological replicate (n = 2); pseudobulk profiles were computed by aggregating single-cell expression data across identified macrophage states. **(B)** Pairwise comparison of TF binding scores between macrophages infected with Salmonella ΔssaV mutant or infected macrophages sorted for SPI2 active WT Salmonella. The volcano plot shows the differential binding score against the −log10(p-value) as provided by TOBIAS. Each point represents a JASPAR motif. All peaks detected in either condition were considered for this analysis. **(C)** UMAP embedding of 2268 genes based on TFBS occupancy profiles. The embedding was generated from the joint analysis of gene-level count matrices of bound TFBS across four conditions (Uninfected, 2hpi, ΔssaV mutant at 18hpi and SPI2 active WT Salmonella at 18hpi). Each point represents one macrophage gene and is positioned according to its TFBS occupancy profile across conditions. Genes are colored according to clusters identified with the Leiden algorithm. Side heatmap shows the mean number of bound TFBS per gene within each cluster across the indicated conditions. **(D)** UMAP embeddings of 2268 genes based on TFBS occupancy profiles as in (C). (Top) Genes are colored according to Stat3 binding at their associated TFBS across conditions, highlighting genes bound for macrophages specifically infected by SPI2 active WT Salmonella (blue), specifically infected by ΔssaV mutant (red), or in both conditions (orange). (Bottom) Genes are colored based on differential expression analysis at the RNA level, indicating genes upregulated in macrophages infected with ΔssaV mutant (orange) or SPI2 active WT Salmonella (green). **(E)** Chord diagrams illustrating the distribution of bound TFBS from five selected TF groups across indicated gene clusters in ΔssaV mutant and SPI2 active WT Salmonella. The width of each connection is proportional to the number of TFBS associated with each TF group–gene cluster pair.

TF footprinting analysis with TOBIAS^30^ revealed transcription factors exhibiting differential binding activity between Δ*ssaV*-infected and SPI2-active WT–infected macrophages (**Figure 5B**). This identified increased activity of AP1 (specifically the TRE motif variant), Maf and Stat families in macrophages harboring SPI2-active WT STm, whereas Nfkb, Cebp, Ets and PARbZip (Nfil3, Dbp, and Tef) families showed the largest binding score increases in Δ*ssaV*-infected BMDMs. We noted two limitations of the previous analysis: (1) motif similarity can obscure precise TF assignment (**Figure S4B**), and (2) even when a TF shows higher overall binding in one condition, many bound TFBS are detected in both conditions (**Figure S4C**). To address these, we refined the footprinting analysis by evaluating TF binding profiles at the level of individual genes across all four conditions (Uninfected, 2 hpi, Δ*ssaV* at 18 hpi, and SPI2-active WT at 18 hpi). Using TOBIAS, we generated a TFBS count matrix (bound TFBS per gene) for each condition and performed a joint analysis to capture dynamic TF binding patterns and identify gene clusters (see **Methods**). To improve TF assignment, only motifs corresponding to TFs expressed at the RNA level were retained; we then focused the analysis on motifs located in chromatin regions uniquely accessible in Δ*ssaV* or SPI2-active WT macrophages. We examined TFBS occupancy for 2,268 genes differentially expressed between Δ*ssaV*-infected and SPI2-active WT–infected macrophages across the four conditions (**Figure 5C**). Unsupervised clustering identified eight gene clusters with similar TFBS profiles. These clusters were partly defined by the distribution of TFBS counts across conditions and were enriched for distinct classes of DEGs (**Figures 5D and S4D**). Consistent with our earlier observations, key transcription factors such as *Stat3* were associated with distinct gene sets depending on condition, rather than exhibiting strictly condition-specific binding (**Figures 5D and S4C**).

We next focused on gene clusters displaying higher TFBS occupancy in Δ*ssaV*-infected (clusters 4 and 5) or SPI2-active WT–infected macrophages (clusters 3 and 6). Notably, these clusters were mirrored across conditions by the core TFs binding their associated genes (**Figure 5E**). Specifically, clusters 4 and 6 were predominantly bound by TFs from the Klf/Sp and Patz1 groups, whereas clusters 5 and 3 were mainly associated with Spi1 (PU.1). These patterns indicate that differential gene expression is associated with, at least in part, a redistribution of TF binding across gene sets. We further observed a marked contribution of AP1 and Maf/Nfe2 TF families in cluster 3, raising the possibility that they contribute toward the anti-inflammatory state. Consistent with this, these motifs, together with Mafb, Maff, Maf, and Stat TFs, were significantly enriched in DEGs upregulated in SPI2-active WT–infected macrophages, whereas Nfkb motifs were preferentially associated with DEGs upregulated in Δ*ssaV*-infected macrophages (**Figure S4E**). Together, these footprinting analyses show that SPI2-associated macrophage reprogramming by STm is accompanied by increased footprint-level engagement of AP1, Maf, and Stat families in the anti-inflammatory program, together with widespread chromatin remodelling involving core macrophage transcription factors.

### CRISPRi perturbations map regulatory circuits underlying macrophage polarization

To test the involvement of the identified TFs and further characterize the regulatory circuitry underlying macrophage polarization, we performed functional perturbation experiments using CRISPR interference. Primary macrophages are refractory to genetic manipulation; we therefore employed the Hoxb8 system as an alternative to BMDMs. This model relies on conditionally immortalized myeloid progenitors that can be expanded over extended periods and induced to differentiate on demand upon removal of β-estradiol from the culture medium^31^. Hoxb8-derived macrophages have been shown to recapitulate key functional properties of primary cells in infection contexts^32^. Importantly, Hoxb8 progenitors are readily amenable to lentiviral transduction, allowing genetic perturbations to be introduced at the progenitor stage prior to differentiation into macrophages^32^.

We selected candidate genes from our scRNA-seq and ATAC-seq analyses and excluded targets not expressed in Hoxb8-derived macrophages or were likely to compromise macrophage viability^33^. We ultimately selected a panel of 87 mouse genes, for which we designed a library of 275 gRNAs (**Figure 6A**). Taking advantage of the flexibility of the Hoxb8 system, we introduced Zim3-dCas9 and the pooled gRNA library into progenitors through two successive lentiviral transductions.

**Figure 6:**
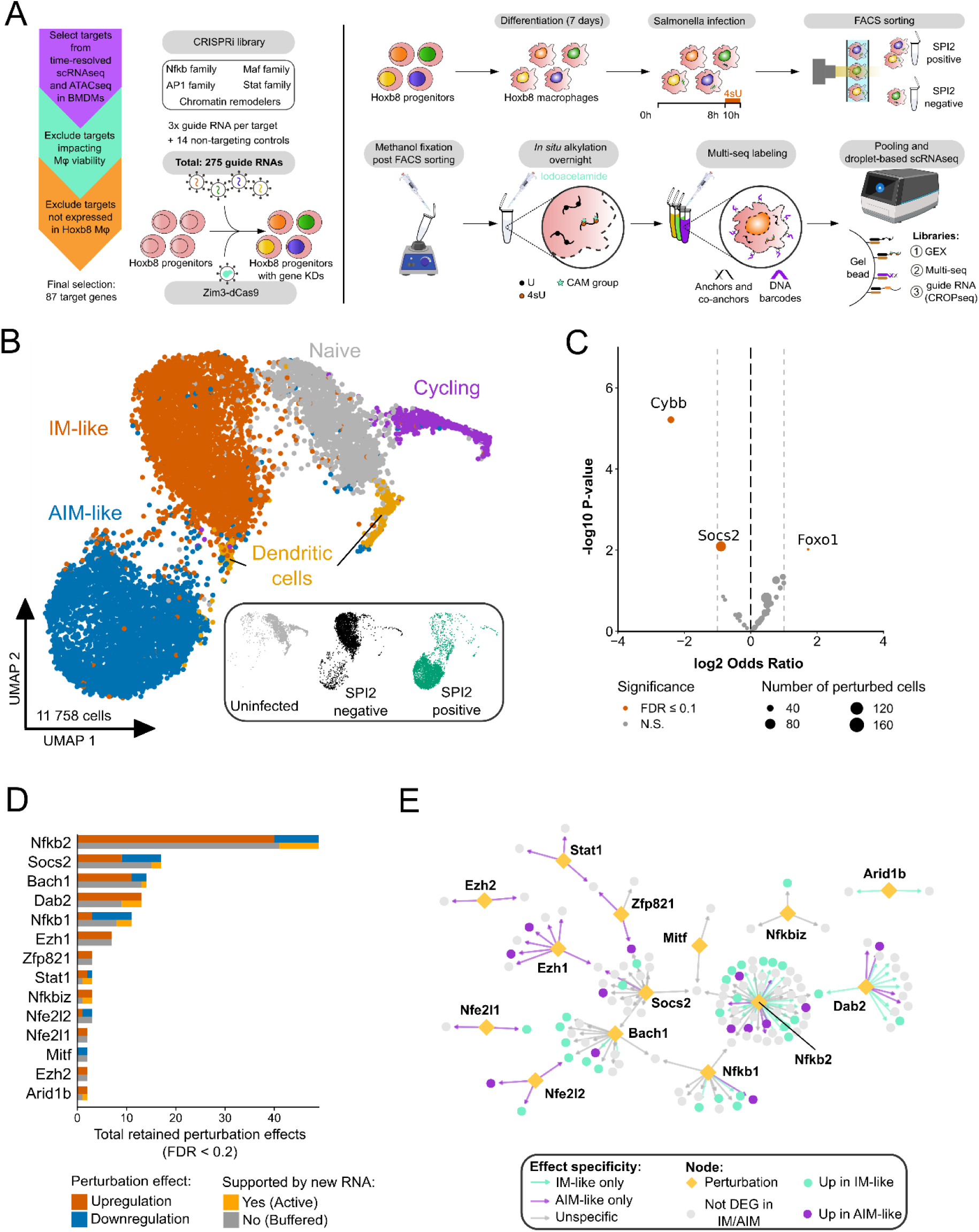
A single-cell CRISPRi screen probes gene networks underlying polarization in Hoxb8 macrophages. **(A)** Experimental workflow for a pooled CRISPRi screening in Hoxb8 macrophages with scRNA-seqscRNAseq readout. (Left) Considerations for designing the guide RNA library and lentiviral transduction into Hoxb8 progenitors. (Right) Workflow of scSLAMseq 2.0, a droplet-based scRNA-seqscRNAseq approach compatible with RNA metabolic labeling, sample multiplexing and capture of guide RNA identity. List of target genes with rationale for their selection and sgRNA sequences are reported in Table S3. **(B)** UMAP embeddings of 11,758 Hoxb8 macrophage transcriptomes obtained with scSLAMseq 2.0. UMAP embeddings result from the joint analysis of new and old RNA levels. Single macrophages are colored by Hoxb8 cell states as identified by unsupervised clustering with the Leiden algorithm. Additionally, the cell repartition from experimental conditions across the dataset are represented as a sub-panel (i.e. Salmonella status based on Multi-seq labeling). **(C)** Volcano plot of differential perturbation abundance between inflammatory and anti-inflammatory Hoxb8 macrophages. The x-axis shows the log2 odds ratio and the y-axis the −log10(p-value). Each point represents a perturbation, with size proportional to the number of cells. Significant perturbation enrichments compared to non-targeting control (NTC) cells are highlighted (FDR < 0.1, binomial generalized linear model with Benjamini–Hochberg correction). **(D)** Summary of sceptre hits based on total RNA levels. Bar heights indicate the total number of significant perturbation effects when targeting the indicated gene (FDR < 0.2, sceptre p-values adjusted with Benjamini–Hochberg correction). Bar colors indicate two additional layers of information: direction of effect (upregulation or downregulation of genes affected by the perturbation) and whether the perturbation effect is also detectable in new RNA levels (Active or Buffered perturbation effects). Only target genes with more than one significant perturbation effect are displayed. **(E)** Gene perturbation network showing significant effects retained in (D). Perturbed genes (diamonds) are connected to their downstream genes (circles). Downstream genes are colored according to their differential expression status (purple: upregulated in anti-inflammatory macrophages; turquoise: upregulated in inflammatory macrophages). Edge colors indicate whether the perturbation effect is detectable specifically in Anti-inflammatory macrophages (purple), Inflammatory macrophages (turquoise) or in multiple macrophage states (grey).

In parallel, we developed scSLAM-seq 2.0, integrating RNA metabolic labelling with sample multiplexing (MULTI-seq^34^) and sgRNA capture (CROP-seq^35^) within a droplet-based scRNA-seq framework, thereby achieving sufficient cell coverage per perturbation (**Figure 6A**). To assess the impact of the CRISPR perturbations on STm infection, transduced progenitors were differentiated into macrophages and infected with the SPI2 reporter strain. Cells were collected at 10 hpi following a 2-hour 4sU labeling pulse, sorted based on SPI2 activity, and methanol-fixed prior to processing with the scSLAM-seq 2.0 workflow.

After sequencing and quality control, we retained 11,758 single-cell transcriptomes, with a median of 1,661 detected genes and 3,920 UMIs per cell (**Figure S5A**). A unique perturbation identity could be assigned to 42% of cells, and 4sU-labeled transcripts were detected at a median of 14.5% of UMIs per cell (**Figures S5B and S5C**). Joint analysis of new and old RNA identified five cell states by unsupervised clustering, consistently recovered across replicates (**Figures 6B and S5D**). Three of these corresponded to macrophage states largely defined by infection condition and STm SPI2 activity. Notably, the two infected states showed transcriptional overlap with the IM *F10* and AIM *Adora3* signatures identified in the BMDM time-course dataset (**Figure S6E**). In addition, we detected a minor population (2.5% of cells) characterized by elevated expression of *Itgax* (*Cd11c*) and *Irf4*, suggestive of a dendritic cell–like phenotype.

To assess the impact of the CRISPR perturbations, we restricted the analysis to macrophages assigned to a single gRNA with demonstrable target repression (3,563 cells; median of 37 cells per targeted gene) (**Figure S5B**). We first asked whether individual perturbations could bias infection outcomes toward inflammatory or anti-inflammatory states. Repression of *Cybb* and *Socs2* skewed cells toward an anti-inflammatory phenotype, whereas repression of *Foxo1* significantly favored an inflammatory state (**Figure 6C**).

Beyond identifying perturbations that markedly influenced infection outcome, we sought to leverage our scRNA-seq readout combined with RNA metabolic labeling to map gene networks underlying macrophage polarization. We applied SCEPTRE^36^ to identify genes whose expression changed in response to each perturbation at the total RNA level. This analysis identified 145 significant perturbation effects (FDR < 0.2), involving 21 targeted genes and 136 downstream genes. Most effects corresponded to upregulation of downstream targets (73%), consistent with factors such as *Nfkb2*, *Bach1*, and *Ezh1* acting predominantly as transcriptional repressors (**Figure 6D**).

We next sought to distinguish perturbation effects that broadly affect the macrophage transcriptional landscape from those that are specific to particular polarization states (**Figure 6E**). Notably, perturbations of *Nfe2l1* and *Nfe2l2* showed detectable effects only in anti-inflammatory macrophages, consistent with the increased activity of the Maf/Nfe2 transcription factor family observed in our TF footprinting analysis. In contrast, a substantial fraction of *Nfkb2*-associated perturbation effects were specific to inflammatory macrophages. Strikingly, perturbations of *Ezh1*/*Ezh2* (PRC2 components) were detectable only in anti-inflammatory states, whereas *Arid1b* (SWI/SNF subunit) showed effects predominantly in inflammatory macrophages, suggesting differential engagement of these chromatin remodeling complexes across polarization states.

Finally, systematic inference of relationships between perturbations and downstream genes does not readily distinguish between direct and indirect effects. Notably, the incorporation of 4sU labelling enabled us to resolve perturbation effects actively occurring at the time of collection and captured in newly synthesized RNA, from effects buffered at the total RNA level (**Figure S5D**). This revealed that most detected perturbation effects are buffered, suggesting that they primarily reflect earlier transcriptional changes during infection rather than ongoing regulatory activity at the time of sampling (**Figures 6D and S5E**).

Taken together, we established a CRISPRi platform suited to probe the transcriptional circuitry of Hoxb8 macrophages during STm infection. Beyond confirming state-specific roles for transcription factors identified by our footprinting analysis, this approach identified two host factors whose perturbation altered infection outcome. Repression of *Foxo1* biased macrophages toward the inflammatory state, implicating FOXO1 in establishment of the anti-inflammatory program. In contrast, repression of *Cybb* shifted cells toward anti-inflammatory macrophages harboring SPI2-active bacteria, consistent with early host antimicrobial activity acting as a checkpoint that determines whether *Salmonella* progresses to the reprogramming stage. Together with the differential engagement of PRC2 and SWI/SNF complexes across states, these results move beyond the STAT3 axis to reveal additional host regulators that shape macrophage fate during infection.

## DISCUSSION

Phagocytosis of *Salmonella* by macrophages initiates a dynamic host–pathogen interplay in which the host cell mounts an inflammatory program aimed at pathogen clearance, while the internalized bacteria deploy countermeasures to withstand this hostile environment and reprogram the host cell toward a more permissive state. In this study, we focused on SPI2-dependent macrophage reprogramming and sought to resolve the temporal sequence of transcriptomic changes underlying this process, linking these dynamics to upstream regulatory TFs.

Our data indicate that infected macrophages initially mount a homogeneous inflammatory response, while reprogramming emerges concomitantly with SPI2 induction around 6 hpi, marked by the activation of anti-inflammatory gene modules. Notably, we observed heterogeneity in this process, with a subset of macrophages rapidly adopting an anti-inflammatory state, whereas others retained an intermediate transcriptional signature up to 8 hpi, suggesting a delayed or gradual reprogramming.

We leveraged time-resolved scRNA-seq combined with RNA metabolic labelling to finely dissect gene expression dynamics. Reconstruction of gene module dynamics along infection trajectories revealed that a substantial fraction of anti-inflammatory genes is already induced as part of the early macrophage response to *Salmonella* infection. These genes are subsequently attenuated in macrophages that control the infection, but reinforced in reprogrammed cells. This pattern is consistent with the idea that SPI2-mediated reprogramming, notably through SteE, hyperactivates an endogenous anti-inflammatory program^14^.

Several studies highlight the central role of the *Salmonella* SPI2 effector SteE in mediating macrophage reprogramming through activation of *Stat3*^26,27,37^. Our regulon- and footprinting-based analyses add a layer to this picture, as *Stat3* was linked to both inflammatory and anti-inflammatory gene modules. This is consistent with STAT3 being activated during infection through more than one route – the SteE/GSK3 axis^26,27^ and SteE-independent inflammatory signalling^28,29^ – dual activity that transcriptional network analyses cannot fully resolve. We also identified several transcription factor groups exhibiting similar binding patterns, including members of the AP-1 family. AP-1 factors have been reported to redistribute across the genome upon stimulation and to recruit chromatin remodelling complexes^38,39^, and multiple epigenetic complexes have been implicated in driving macrophage activation^40,41^. Together, these observations are consistent with widespread chromatin remodelling accompanying macrophage reprogramming.

Unexpectedly, *Stat3* knockdown in Hoxb8 macrophages did not measurably affect infection outcome. Notably, a recent study reported that *Salmonella* infection elicits only a weak anti-inflammatory response in Hoxb8 macrophages, which lack several canonical M2 markers^42^. This points to important differences in the polarization circuitry of Hoxb8 macrophages compared to primary BMDMs, and therefore perturbation phenotypes in this system warrant confirmation in primary cells. By contrast, Foxo1 knockdown increases the proportion of inflammatory Hoxb8 macrophages able to control infection, indicating that factors beyond STAT3 contribute to macrophage reprogramming. Despite targeting 14 of 87 AP-1–related genes, we did not detect a prominent effect on infection outcome or perturbation phenotypes. This may reflect functional redundancy among AP-1 family members, or the limited number of cells recovered per perturbation.

While our study delineates transcriptional programs that bias macrophage polarization toward distinct states, a key outstanding question is what determines whether *Salmonella* successfully initiates host cell reprogramming. Notably, knockdown of *Cybb* (gp91 phox) emerged as the most prominent hit in our CRISPR screen, biasing infection toward anti-inflammatory macrophages harbouring SPI2-active bacteria. Because our time-resolved data place the onset of reprogramming at 4-6 hpi, there is an early window before reprogramming takes hold during which the host can still restrict the bacteria; that *Cybb*, encoding a core component of the NADPH oxidase complex, is the strongest hit suggest host antimicrobial capacity in this window is a key determinant of whether *Salmonella* reaches the reprogramming stage. We note this rests on a single perturbation, and NADPH-oxidase-derived ROS could also influence polarization independently of bacterial killing; distinguishing these will require targeted follow-up. Future work could also leverage single SPI2 effector perturbations to investigate their relative contribution to the modulation and reprogramming of macrophages.

## SUPPLEMENTARY TABLES

**Table S1: Metadata of 578 single-cell libraries included in analysis - related to Figures 1, 2, 3 and 4**

**Table S2: Differentially expressed genes from modules linked to Inflammatory and Anti-inflammatory macrophages - related to Figure 3**

**Table S3: Summary of selected target genes and sgRNA sequences used for single-cell CRISPRi screen in Hoxb8 macrophages - related to Figure 6**

**Table S4: *Salmonella* strains, plasmids and DNA oligos used in this study**

## MATERIAL & METHODS

## EXPERIMENTAL PROCEDURES

### Cell culture

#### Salmonella strains

*Salmonella enterica* serovar Typhimurium strain SL1344 was used across the present study. Activity of the *ssaG* promoter was detected with the pFCcGssaG reporter plasmid. This plasmid was constructed by […]. The Δ*ssaV* mutant strain was a gift from Andrew Grant^43^. See Table S4 for details about all strains, plasmids and primers.

### Bone marrow derived macrophages

Primary bone marrow derived macrophages (BMDMs) were produced from bone marrow extracted from the femur of 6 to 8-week old female C57BL/6 mice (Charles River). Red blood cells were lysed in 0.83% NH_4_Cl for 5 minutes. The remaining progenitor cells were cultured at 37°C and 5% CO_2_ in Dulbecco’s modified eagle medium with high glucose (DMEM; Sigma) containing 20% L929 (ATCC #CCL-1) conditioned media (LCM), 10% fetal bovine serum (FBS; Gibco), 10 mM HEPES (Sigma), 1 mM sodium pyruvate (Sigma), 0.05 mM beta-mercaptoethanol (Sigma), 100 U/mL penicillin (Sigma), and 100 μg/mL streptomycin (Sigma). Fresh medium was supplemented on day 3. Differentiated BMDMs were harvested at day 7 and seeded in DMEM without LCM or antibiotics (BMDM infection medium) for infection.

### Cell lines

#### Hoxb8

Hoxb8 progenitors were maintained in RPMI 1640 medium (Gibco, #21870-076) supplemented with 10% FBS, 2 mM L-glutamine (Gibco, #25030-024), 100 U/mL penicillin, 100 µg/mL streptomycin (Gibco, #15140-122), 20 ng/mL GM-CSF (Peprotech) and 1 µM β-estradiol (Sigma, #E2758) and cultured at 37 °C and 5% CO_2_. Progenitors were maintained in 6-well plates (Sigma, #CLS3516) and split every other day to maintain low density or expanded in T-75 flasks (Sarstedt, #83.3911.002).

To differentiate Hoxb8 progenitors into macrophages for *Salmonella* infection, cells were washed twice with DPBS (Gibco, #14190-144) to remove β-estradiol and resuspended into differentiation medium made of: RPMI 1640 medium supplemented with 10% FBS, 2mM L-glutamine, 100 U/mL penicillin, 100 μg/mL streptomycin, 20 ng/mL GM-CSF (Peprotech) and 20 ng/mL M-CSF (Peprotech). Cells were seeded into 6-well plates at 350, 000 cells/well in 2 mL of differentiation medium. Differentiating cells were given an additional 1 mL of differentiation medium on day 3 and day 5 after initiating differentiation. On day 7, cells were switched to Hoxb8 differentiation medium without antibiotics (Hoxb8 infection medium) after washing with 2 mL of pre-warmed DPBS. Differentiated Hoxb8 macrophages were used for *Salmonella* infection on day 8.

#### HEK293T

HEK293T cells were maintained in high glucose DMEM medium (Gibco, #11960-044) supplemented with 10% FBS, 2 mM L-glutamine, 100 U/mL penicillin and 100 µg/mL streptomycin and cultured at 37 °C and 5% CO_2_.

#### Mycoplasma contamination assay

Cell lines were routinely controlled for Mycoplasma contamination by qPCR (Mycoplasmacheck, Eurofins).

### scSLAM-seq

#### *In vitro* infection of BMDMs, RNA metabolic labeling and single-cell sorting

Bacteria were grown in MgMES minimal medium, pH 5.0 (170 mM 2-(N-morpholino)ethanesulfonic acid (MES), 5 mM KCl, 7.5 mM (NH_4_)_2_SO_4_ , 0.5 mM K_2_SO_4_ , 1 mM KH_2_PO_4_ , 8 mM MgCl_2_ , 38 mM glycerol, and 0.1% casamino acids) for 16 hours. Bacteria were opsonized with 8.5% mouse serum (i.e. 45 µL of *Salmonella* culture, 20 µL of mouse serum and 170 µL of BMDM infection medium) for 20 minutes at room temperature followed by a 1:3.5 dilution with BMDM infection medium. Opsonized bacteria were added to BMDMs at MOI 5-10, resulting in an uptake of one bacterium per infected macrophage. Infection was synchronized by 5 minutes of centrifugation at 110g followed by an incubation for 25 minutes at 37°C with 5% CO_2_ to allow for phagocytosis. Infected BMDMs were washed 3x with pre-warmed DPBS and incubated in infection medium supplemented with 20 µg/mL gentamicin for the remainder of the infection. Two hours prior to harvesting, 4-thiouridine (4sU) in water was added to a final concentration of 150 µM. At desired time points, BMDMs were washed 3x with DPBS and finally harvested in ice-cold DPBS by gentle scrapping. Collected macrophages were filtered to remove cell clumps and run on a BD FACS Aria III sorter. Apoptotic cells, doublets and uninfected macrophages were excluded by gating. Infected BMDMs were sorted with continuous cooling at 4°C in 96-well plates pre-filled with 4 µL of lysis buffer (2 U/µL of RNAse inhibitor and 0.2% Triton X-100 in nuclease-free water). Fluorescence parameters associated with each sorted single cell were recorded by index sorting. Plates with sorted cells were immediately sealed, briefly spun down and stored at - 80°C.

#### scSLAM-seq cDNA library preparation

Single-cell SLAM-seq libraries were prepared following our previous protocol^18^. Plates with sorted single-cells were thawed at room temperature for 3 minutes. Alkylation of 4sU was set up by adding 0.4 µL of 10X PBS to each well followed by 4.4 µL of 20 mM iodoacetamide in DMSO. Reaction was incubated at 50°C for 5 minutes in a thermocycler. Alkylation was quenched by adding 1.3 µL of 0.1M DTT and incubation at room temperature for 5 minutes. ERCC Spike-in (Invitrogen, #4456740) controls were added to each well (0.3 µL of a 1:2 million dilution). Reaction clean-up was performed with 10 µL of RNA XP beads with an incubation time of 10 minutes at room temperature for RNA capture followed by washing with 50 µL of fresh 80% EtOH for a total of two washings. Bead pellets were dried for a maximum of 2 minutes. The SMART-Seq v4 Ultra Low Input RNA kit (Takara) was then used to perform the reverse transcription using one quarter of the recommended reagent volumes. Dried bead pellets were resuspended in 4 µL of CDS buffer (0.2 µL of RNAse inhibitor, 0.5 µL of CDS primer and 3.3 µL of nuclease-free water per well). Then, plates were centrifuged to pellet beads, incubated at 72°C for 3 minutes and cooled down at 4°C for 3 minutes in a thermocycler. Immediately, 1.9 µL of 1st strand synthesis master mix (1 µL of 5X Ultra-low first strand buffer, 0.25 µL of SMART-seq v4 Oligo, 0.125 µL of RNAse inhibitor and 0.5 µL of SMARTScribe Reverse Transcriptase per well) was added. Plates were centrifuged to pellet beads and incubated at 42°C for 90 minutes and 70°C for 10 minutes in a thermocycler. For second strand synthesis and cDNA amplification 7.5 µL of master mix (6.25 µL of 2X SeqAmp PCR Buffer, 0.25 µL of PCR Primer II A, 0.25 µL of SeqAmp DNA Polymerase and 0.75 µL of nuclease free water per well) were added. Plates were centrifuged to pellet beads and reaction was run in a thermocycler with the following program: (1) initial denaturation at 95°C for 1 minute, (2) 21x cycles of 98°C for 10s / 65°C for 30s / 68°C for 3 minutes, and (3) final incubation at 72°C for 10 minutes. cDNA products were cleaned with 12.75 µL of AMPure XP beads (Beckman, #A63881) with an incubation time of 8 minutes at room temperature for cDNA capture followed by washing with 100 µL of fresh 80% EtOH for a total of two washings. Bead pellets were dried for a maximum of 3 minutes. Dried pellets were resuspended in 17 µL of elution buffer and left at room temperature for 5 minutes. Finally, 14 µL of clear supernatant was saved after bead separation using a magnet. Quality control of libraries was done with cDNA quantification using Qubit 3.0 Fluorometer with the dsDNA High sensitivity assay (Invitrogen, #Q32851), and cDNA size distribution profile was checked using a 2100 Bioanalyzer with the High Sensitivity DNA kit (Agilent, #5067-4626).

#### scSLAM-seq cDNA library tagmentation and sequencing

Libraries were tagmented using the Nextera XT DNA Library Preparation Kit (Illumina, #FC-131-1096) with one quarter of the recommended reagent volumes. Starting input was 0.5 ng of cDNA (1.25 µL of a 0.4 ng/uL dilution). Tagmentation reaction was set up by adding 3.75 µL of master mix made of 2.5 µL of tagmentation DNA buffer and 1.25 µL of amplicon tagment mix buffer per well, followed by an incubation at 55°C for 10 minutes and a cooling down at 10°C in a thermocycler. Then, 1.25 µL of neutralizing buffer was immediately added to each well and plates were left at room temperature for 5 minutes. For library amplification and indexing, unique combinations of i7 and i5 indexes were chosen for each library (1.25 µL of each index was added per well) and 3.75 µL of Nextera PCR master mix was dispensed to each well. Reactions were run in a thermocycler with the following program: (1) 72°C for 3 minutes, (2) 95°C for 30s, (3) 13x cycles of 95°C for 10s / 55°C for 30s / 72°C for 1 minute and (4) 72°C for 5 minutes. PCR products were cleaned with 7.5 µL of AMPure XP beads with an incubation time of 10 minutes at room temperature for cDNA capture followed by washing with 100 µL of fresh 80% EtOH for a total of two washings. Bead pellets were dried for a maximum of 3 minutes. Dried pellets were resuspended in 13 µL of resuspension buffer and left for elution at room temperature for 5 minutes. Finally, 10 µL of clear supernatant was saved after bead separation using a magnet. For quality control, cDNA was quantified by Qubit 3.0 Fluorometer with the dsDNA High sensitivity assay (ThermoFisher) and library profiles were checked using a 2100 Bioanalyzer with High Sensitivity DNA kit (Agilent).

Libraries were pooled together at equimolar ratios for sequencing, aiming at 5 millions reads per library. Sequencing was done as 4 separate runs on a NovaSeq6000 sequencer using SP/S1 flow cells in paired-end mode with 101x cycles for each read.

### Paired ATAC-seq and RNA-seq

#### Integrated Assay for Transposase Accessible Chromatin (ATAC) Sequencing and RNA Sequencing

BMDMs were infected with the *Salmonella* SPI2 reporter strain (WT or ΔssaV mutant) as described above (without exposure to 4-thiouridine for RNA metabolic labeling), collected and gated for Fluorescence Activated Cell Sorting (FACS).

To generate ATACseq libraries, we sorted 50,000 macrophages containing *Salmonella* (i.e. mCherry+) by FACS per sample. For uninfected macrophage samples, 50,000 macrophages were isolated by FACS without consideration of mCherry-status. Following isolation of the macrophages, ATAC-Seq libraries were immediately generated using the Omni-ATAC protocol^44^. Libraries were sequenced on an Illumina NextSeq 2000 sequencer using paired-end sequencing with 101x cycles for each read (average of 31M reads per library).

For RNA-seq on matched samples, we sorted 50,000-100,000 infected macrophages containing *Salmonella* (i.e. mCherry^+^) by FACS per sample. For uninfected macrophage samples, 50,000-100,000 macrophages were isolated by FACS without consideration of mCherry-status. Sorted macrophages were then centrifuged at 500 x *g* for 5 min at 4 °C, the supernatant was removed, and macrophage pellets were snap frozen in liquid nitrogen and stored at -80 °C until RNA was isolated. RNA was isolated using the Quick DNA-RNA Miniprep Kit (Zymo Research) following the manufacturer’s protocol. Following RNA isolation, mRNA-Seq libraries were generated using the NEBNext Ultra II Directional RNA Library Kit for Illumina (NEB E7765), the NEBNext Poly(A) mRNA Magnetic Isolation Module (NEB E7490), and the NEBNext Multiplex Oligos for Illumina (NEB E7335/E7500/E7710) following the manufacturer’s protocol. Libraries were sequenced on an Illumina NextSeq 2000 sequencer using single-end sequencing with 76x cycles per read (average of 45.5M reads per library).

### scCRISPR / CROP-seq

#### Construction of CROP-seq-opti-Puro-T2A-TagRFP657

Similarly to a previous study^20^, we modified the CROP-seq-opti^45^ vector (Addgene #106280) to facilitate lentivirus titration by adding a T2A-TagRFP657 cassette downstream of the puromycin resistance marker.

Briefly, PCR fragments of CROPseq-opti amplified with Q5 High-Fidelity 2X Master Mix (NEB, #M0492S) and the T2A-TagRFP657 insert (IDT, gBlock) were assembled by Gibson assembly (NEB, #E2621S) followed by transformation into NEB stable competent cells (NEB, #C3040H). Clones were screened by colony PCR and validated by whole-plasmid sequencing prior to plasmid preparation using the ZymoPURE II Plasmid Midiprep Kit (Zymo Research, #D4200). All primers used for CROP-seq-opti-Puro-T2A-TagRFP657 construction are listed in Table S4. To prepare the vector for sgRNA library preparation, we digested CROP-seq-opti-Puro-T2A-TagRFP657 with BsmBI (NEB, #R0739S), followed by dephosphorylation with Shrimp Alkaline Phosphatase (NEB, #M0371S) and agarose-gel extraction of the digested product (Macherey-Nagel, #740609.50).

#### Construction of the pooled sgRNA library

Target genes were selected based on the following criteria: (1) the target gene is a hit from our scSLAM-seq or ATAC-seq analysis, (2) its knockout does not significantly impact cell viability according to a previous CRISPR screen in immortalized bone-marrow-derived macrophages^33^, and (3) the target gene is expressed in both primary BMDMs and differentiated Hoxb8 macrophages (in-house scRNA-seq dataset). We also picked additional targets from the literature (e.g. chromatin remodeler complexes) that were related to our hits. All target genes and sgRNA sequences are listed in Table S3.

sgRNA sequences targeting selected genes were designed with CRISPick^46^ (CRISPRi for SpyoCas9 and RS3i on-target scorer) to obtain 261 sgRNAs targeting 87 genes (three sgRNAs per gene) and 14 non-targeting controls (NTCs). Finally, sgRNAs were ordered as a single-stranded DNA oligo pool (IDT, oPool) with the following format:

5’-GGCTTTATATATCTTGTGGAAAGGACGAAACACCG [20bp sgRNA sequence] GTTTAAGAGCTATGCTGGAAACAGCATAGCAAGTT-3’

The oligo pool was inserted into BsmBI-digested CROP-seq-opti-Puro-T2A-TagRFP657 by Gibson assembly : 75 ng (13 fmol) of plasmid vector were mixed with 72 ng (2.582 pmol) of single-stranded DNA oligo pool (resuspended to 100 ng/uL), 10μl of HiFi DNA Assembly Master Mix (NEB, #E2621S) and nuclease-free water up to 20 µL (Invitrogen, #AM9932). A control reaction without insert was set up in parallel. The reactions were incubated at 50°C for 60 minutes. Gibson products were cleaned up with 15 µL of AMPure XP beads with an incubation time of 15 minutes at room temperature for capture followed by washing with 100 µL of fresh 80% EtOH for a total of two washings. Bead pellets were dried for a maximum of 3 minutes. Dried pellets were resuspended in 3 µL of nuclease-free water and left for elution at room temperature for 10 minutes. Next, 2 µL of purified Gibson product was electroporated into 20 µL of ElectroMAX Stbl4 Competent Cells (Invitrogen, #11635018) with a Gene Pulser Xcell system (Bio-Rad, #1652660) and the following parameters: 1.8 kV, 25 µF, 200 Ohm. After recovery, 50 µL of cells electroporated with the sgRNA library and 50 µL of cells electroporated with control Gibson were plated on standard LB agar plates (100-mm Petri dish, carbenicillin) to estimate library coverage. Remaining cells electroporated with the sgRNA library were expanded in liquid culture (50 mL LB medium, carbenicillin) and were grown at 28°C, 225 rpm for 24 h. The sgRNA library was extracted from the liquid culture using the ZymoPURE II Plasmid Midiprep Kit (Zymo Research, #D4200) with removal of endotoxins.

Quality control of the sgRNA library for sgRNA insertion and representation was performed by next-generation sequencing. Briefly, 20 ng of extracted sgRNA library was amplified by PCR with 2.5 µL of a 10 µM forward/reverse primer pair (CT066 and CT016, see Table S4), 25 µL of NEBNext High-Fidelity 2X PCR Master Mix (NEB, #M0541S) and nuclease-free water up to 50 µL (Invitrogen, #AM9932). The reaction was run in a thermocycler with the following program: (1) 98°C for 30 seconds, (2) 23x cycles of 98°C for 10 seconds / 66°C for 30 seconds / 72°C for 30 seconds and (3) 72°C for 2 minutes. PCR products were cleaned with DNA Clean & Concentrator-5 kit (Zymo Research, #D4003). Purified PCR products were sequenced by Nanopore sequencing.

#### Lentivirus production and titration

Low-passage HEK293T cells at 70% confluency in T-25 flasks were carefully washed with pre-warmed DPBS and fed 4 mL of fresh medium. Next, cells were co-transfected with 1.5 µg of psPAX2 (Addgene, #12260), 1 µg of pMD2.G (Addgene, #12259) and 2.5 µg of either pHR-UCOE-EF1a-Zim3-dCas9-P2A-BFP (Addgene, #188777) or our sgRNA plasmid library using Fugene HD transfection reagent (Promega, #E2311). Transfected cells were incubated at 37 C° and 5% CO_2_ for 48 hours, then supernatants were collected and centrifuged at 200g, 4 °C for 5 minutes. Cleared supernatants were filtered through 0.45 µm filters (Sarstedt, #83.1826) and frozen at -80 °C as single-use 1 mL aliquots.

Lentivirus stocks of our sgRNA library were titrated by transducing Hoxb8 progenitors (as described below). Then, progenitors were collected at 48 hours post-transduction, fixed with 4% paraformaldehyde (Sigma, #158127) for 15 minutes at room temperature and washed with DPBS. Fixed cells were then run on a flow cytometer (Agilent, NovoCyte Quanteon) to determine the percentage of TagRFP657 positive cells.

#### Lentiviral transductions of Hoxb8 progenitors

The lentiviral transduction tailored for Hoxb8 progenitors was adapted from a published protocol^47^. We delivered dCas9 and the sgRNA library in two successive transductions.

For transduction of dCas9, progenitors were seeded into 12-well plates (100, 000 cells per well in 0.5 mL of medium). The next day, polybrene (Sigma, #TR-1003) was added to a final concentration of 5 µg/mL, followed by 200 µL of lentivirus and cells were spinoculated at 1000g, 30 °C for 1 hour. Following spinfection, 2.3 mL of pre-warmed medium was added and cells were incubated at 37 °C and 5% CO_2_ overnight. The next day, 2 mL of medium were slowly removed, followed by the addition of 1 mL of pre-warmed medium. Progenitors were maintained in culture and splitted as necessary. At 7 days post-transduction cells were collected, washed with pre-warmed DPBS and run on a BD FACS Aria III sorter. Apoptotic cells and doublets were excluded by gating followed by sorting at room temperature for BFP positive progenitors.

For transduction of the sgRNA library, dCas9-expressing (BFP-positive) Hoxb8 progenitors were transduced using the same protocol, with titrated lentiviral input adjusted to achieve a multiplicity of infection (MOI) of 0.1 to ensure delivery of 1 sgRNA per progenitor. Cell numbers were also scaled to achieve a coverage of 1,000 transduced cells per sgRNA. At 48 hours post-transduction, cells were selected with 2 µg/mL puromycin (Sigma, #P4512) for 72 hours, until complete cell death was observed in a non-transduced control well. Selected Hoxb8 progenitors were maintained in culture, passaged and expanded while preserving a minimal 1,000x cell coverage per sgRNA. At 7 days post-transduction, proper selection (TagRFP657 positive cells) was assessed by flow cytometry. If necessary, the cell population was FACS-sorted again to re-enrich for BFP-positive cells while maintaining library coverage. Progenitors were then allowed to recover for 72 hours before initiating macrophage differentiation as described earlier (see Cell Culture section).

#### *In vitro* infection of Hoxb8 macrophages, RNA metabolic labeling and bulk sorting

*Salmonella* pFCcGssaG culture and infection of Hoxb8 macrophages were performed with the same protocol and conditions as for BMDMs (except that Hoxb8 infection medium was used for bacterial opsonization), up to the addition of gentamicin-containing medium. Two hours prior to harvesting, 4sU was added to a final concentration of 200 µM. At 10 hours post-infection, Hoxb8 macrophages were washed three times with DPBS and harvested in ice-cold DPBS by gentle scraping. Collected macrophages were filtered to remove cell clumps and run on a BD FACS Aria III sorter. Apoptotic cells, doublets and uninfected macrophages were excluded by gating. We sorted 500,000 - 1,000,000 infected macrophages with SPI2 active or SPI2 inactive intracellular *Salmonella* in separate 15 mL tubes with continuous cooling at 4°C.

Sorted cells were centrifuged (300 × g, 4°C, 5 minutes) and resuspended as single-cell suspensions in 200 µL ice-cold DPBS. Cells were fixed by dropwise addition of 800 µL pre-chilled methanol (−20°C) with gentle vortexing. Following incubation on ice for 20 min, with gentle mixing every 5 min, fixed cells were stored at −80°C.

#### scSLAMseq 2.0: *In situ* alkylation, cell rehydration and MULTI-seq labeling

For alkylation, a 8.4 mM solution of iodoacetamide (IAA) was freshly prepared by resuspending 9.3 mg of IAA (Thermo Scientific, #A39271) into 1.2 mL of DPBS and adding 4.8 mL of chilled methanol. Methanol-fixed cells were thawed on ice for 15 minutes before centrifugation at 1000 x g, 4°C, for 5 minutes. Cell pellets were resuspended into 1 mL of 8.4 mM IAA and incubated at 4°C for 18 hours, protected from light.

For rehydration, alkylated cells were centrifuged at 800 x g, 4°C, for 5 minutes to remove methanol and IAA. Cell pellets were resuspended into 500 µL of 3X SSC + 0.04% BSA buffer (SSC: Invitrogen, #AM9763; BSA: Invitrogen, #AM2616) and incubated on ice for 20 minutes.

For sample multiplexing using MULTI-seq labeling^34^, rehydrated cells were centrifuged (400 x g, 4°C, for 5 minutes), washed with 500 µL of 3X SSC buffer without BSA and finally resuspended in 180 µL of the same buffer. Lignoceric acid anchor (Sigma, #LMO001-30RXN) and sample-specific DNA barcodes^34^ were mixed in equimolar ratios (4 µM each) in 1× DPBS, and 20 µL of this mixture was added to each cell sample, followed by incubation on ice for 5 min. Subsequently, 20 µL of 4 µM palmitic acid co-anchor (Sigma, #LMO001-30RXN) was added, and samples were incubated on ice for an additional 5 min. Cells were washed twice with 500 µL of ice-cold 3X SSC + 1% BSA buffer and finally resuspended into 50 µL of ice-cold 0.125X SSC + 0.04% BSA buffer.

Cell samples were counted and combined to achieve the desired representation in the final cell pool. The pooled suspension was filtered with a 40 µm cell strainer (pluriSelect, #43-10040-50) and kept on ice until loading onto the microfluidic chip for droplet-based scRNA-seq.

#### scSLAMseq 2.0: droplet-based scRNA-seq, library preparation and sequencing

The pooled cell suspension was loaded onto a Chromium X Controller (10x Genomics, #1000331) across multiple lanes to achieve the desired number of recovered cells. Samples were processed using the Chromium GEM-X Universal 3′ Gene Expression v4 kit (10x Genomics, #1000686) according to the manufacturer’s protocol for droplet generation, reverse transcription, cDNA amplification, and the gene expression (GEX) library construction, with the following modifications: (1) in Step 2.2.a, 1 µL of 2.5 µM MULTI-seq primer^34^ was added to the cDNA amplification master mix; and (2) in Step 2.3.d, the supernatant was saved, as it contains the MULTI-seq barcodes.

For the MULTI-seq libraries, 80 µL of saved supernatant containing MULTI-seq barcodes was mixed with 260 µL of SPRIselect beads (Beckman, #B23318) and 180 µL of isopropanol (Sigma, #33539). After 5 minutes of incubation, the bead pellets were washed twice with 500 µL of 80% EtOH. After air-drying, bead pellets were resuspended into 50 µL of EB buffer (Qiagen, #1014609). Next, the library was prepared by PCR: 3.5 ng of eluted cDNA barcodes were mixed with 2.5 µL of Universal I5 primer (10 µM), 2.5 µL of RPI primer (10 µM), 26.25 µL of Kapa HiFi HotStart ReadyMix (Roche, #KK2601) and nuclease-free water up to 50 µL. PCR reactions were run in a thermocycler with the following program: (1) 95°C for 5 minutes, (2) 8-12x cycles of 98°C for 15 seconds / 60°C for 30 seconds / 72°C for 30 seconds and (3) 72°C for 1 minute. PCR products were cleaned with 80 µL of SPRI beads, 2 washings with 200 µL of 80% EtOH and a final resuspension with 25 µL of EB buffer.

For the CROP-seq libraries, we performed three rounds of PCRs starting from the same full-length cDNA used to generate the GEX libraries (Step 2.3.m), as previously described^45,48^. For the first PCR, 10 ng of full-length cDNA was combined with 3 µL of a 10 µM forward/reverse primer pair (CT067 and CT068, see Table S4), 5 µL of KAPA SYBR FAST qPCR Master Mix (Roche, #KK4600), and nuclease-free water to a final volume of 10 µL. Reactions were performed using the following cycling conditions: (1) initial denaturation at 95°C for 5 minutes , followed by (2) amplification cycles of 95°C for 30 seconds and 65°C for 45 seconds. Amplification was monitored in real time and stopped prior to reaching the plateau phase. PCR products were purified using 20 µL AMPure XP beads after dilution with 10 µL nuclease-free water, followed by two washes with 80% ethanol and final elution in 20 µL nuclease-free water. The second PCR was performed using the same conditions, with 1 µL of a 1:25 dilution of the purified first PCR products as input and a new primer pair (CT069 and CT070, see Table S4). The third PCR was carried out similarly using 1 µL of a 1:25 dilution of the purified second PCR products, with 1.5 µL of 10 µM CT070 primer and 1.5 µL of Nextera XT i7 index primer (Illumina, #FC-131-2001). In addition, the annealing/extension step was performed at 72°C instead of 65°C. Quality of libraries was assessed by quantification using a Qubit 3.0 Fluorometer with the dsDNA High Sensitivity assay (Thermo Fisher Scientific), and library size profile was evaluated using a 2100 Bioanalyzer with the High Sensitivity DNA kit (Agilent).

Libraries were pooled together aiming at sequencing depths of 90,000 reads per cell for GEX libraries and 5,000 reads per cell for both MULTI-seq and CROP-seq libraries. Sequencing was performed on a NovaSeq X Plus platform using a 10B flow cell in paired-end mode (Read 1: 28 cycles; Read 2: 90 cycles; Index 1: 10 cycles; Index 2: 10 cycles).

## BIOINFORMATIC ANALYSIS

### scSLAM-seq data analyis

#### Flow cytometry and Index sorting data analysis

Flow cytometry data were analysed at the population level with CytoExploreR (v1.1). Index sorting data were read from fcs files with flowCore (v1.11.2) to extract fluorescence measurements of sorted single cells.

#### Preprocessing of scSLAM-seq data

Raw fastq files were generated by demultiplexing with bcl2fastq and general quality control was performed with FastQC v0.12.1 and MultiQC v1.15. Sequencing reads were trimmed with Cutadapt v4.4 to remove the first 10bp of each read, the Nextera XT adapters (CTGTCTCTTATACACATCT), poly-A tails and low quality bases (Phred score < 20). Reads shorter than 20bp after trimming were excluded. Trimmed reads were aligned to a combined reference of mouse (Ensembl v105) and *Salmonella* Typhimurium SL1344 (ASM21085v2) genomes as well as ERCC spike-ins. Alignment was done with STAR v020201 (and the following additional parameters: --outFilterMismatchNmax 20; --outFilterScoreMinOverLread 0.4; --outFilterMatchNminOverLread 0.4; --alignEndsType EndToEnd; --outSAMattributes nM MD NH). Bam files were converted to CIT files and merged together with the GEDI toolkit^49^. Mismatch analysis, estimation of new-to-total RNA ratios (NTRs) per gene across cells and generation of the read count matrix was done with GRAND-SLAM v2.0.6^22^ (and the following additional parameters: -trim5p 15 -minEstimateReads 7500 -allGenes).

#### Selection of genes consistently detected with precise NTR estimates

*Salmonella* genes were excluded from selection due to sparse detection across cells and the predominance of rRNA transcripts.

Mouse genes reliably detected across cells were identified using external RNA spike-ins (ERCC). For each cell, the relationship between known ERCC input molecule numbers and observed read counts was modeled using LOESS regression. Cells with poor concordance between expected and observed ERCC levels (Spearman correlation < 0.7) were excluded. Detection thresholds derived from ERCC-based models were then used to define gene expression sensitivity per cell, and genes detected above this threshold in at least 5% of cells were retained.

To ensure robust estimation of new RNA levels, genes with unreliable new-to-total RNA ratio (NTR) estimates were filtered out. For each gene, the width of the credible interval associated with NTR estimates (as computed by GRAND-SLAM) was quantified across expressing cells. Genes with high uncertainty relative to their estimated NTR (credible interval to NTR ratio > 1) were excluded.

#### Cell quality control and normalization of count matrix

A metadata table was built based on various metrics and measurements collected for each cell from the index sorting data, the mapping results and GRAND-SLAM analysis (see Table S1). The total RNA count matrix was reduced to selected mouse genes (see previous section) and cells with less than 2,500 detected genes, more than 3% mitochondrial reads, more than 3% ERCC reads or less than 25% exonic reads were excluded.

The filtered total RNA count matrix was then normalized with SCnorm v1.12.1^50^ after removal of *Salmonella* and ERCC counts. Finally, the count matrices for new and old RNA were computed based on the normalized total RNA count matrix and NTR matrix.

#### Dimensional reduction and clustering based on new and old RNAs

Analysis was conducted in R with the Seurat v4.0 package. For joint analysis of new and old RNAs, we included in the analysis only cells exposed to 4sU with sufficient RNA labeling overall (578 cells). Both new and old RNA matrices were analysed in parallel to log-transform the normalized counts, identify the top highly variable genes (top 1,000 for old RNA and top 500 for new RNA) and perform a principal component analysis (irlba v2.3.5.1). Then, a joint PCA matrix was built by combining the top 4 PCs from each assay (new and old) before dimensional reduction with UMAP. The joint PCA was also used to build a KNN graph (k = 10) followed by cell clustering with the Leiden algorithm^51^. We used the C implementation for R of the Leiden algorithm (leidenbase v0.1.3, Trapnell Lab) with a resolution set at 0.07.

#### Dimensional reduction based on total RNA

To generate the UMAP related to Figure S1D, a scRNA-seq analysis focused on total RNA levels was performed including all single-cells passing quality control, regardless of their RNA labeling status (765 cells). Similarly to the joint analysis, the top 1,000 highly variable genes were selected and used to perform PCA. The top 6 PCs were used for dimensional reduction with UMAP.

#### Differentially expressed gene testing and pathway enrichment analysis

For differential expression gene (DEG) testing, we generally used the built-in Seurat function *FindMarkers()* with a Wilcoxon Rank Sum test followed by p-value adjustment for multiple comparisons with the Benjamini-Hochberg procedure to obtain a False Discovery Rate (FDR). Otherwise stated, we performed DEG testing on new RNA levels and used FDR < 0.05 as a significance cut-off. To compute gene module scores, we used the built-in Seurat function *AddModuleScore()* with default parameters and new RNA levels.

For pathway enrichment analysis, gene ontology (GO) enrichment was conducted on selected gene sets using EnrichR(v3.1) and querying the GO Biological Process 2025 database. Enrichment significance was evaluated using Fisher’s exact test with Benjamini–Hochberg correction for multiple comparisons. GO terms with an adjusted p-value < 0.05 were considered significant. For visualization (Figure 2D), the top 50 terms per gene set, ranked by adjusted p-value, were selected.

#### Gene module definition and correlation with *Salmonella* SPI2 activity

Gene modules in Figure 3A were defined using two differential expression analyses: (1) AIM *Adora3* versus IM *F10* and Intermediate state, and (2) IM *F10* versus AIM *Adora3* and Intermediate state. Genes were assigned to modules based on the direction of the average log_2_ fold change (positive: modules A and D; negative: modules C and B). In cases of overlap, >modules A and D were prioritized. Genes within a module were further classified as early or delayed induction based on differential expression in IM *Tnfsf9* (2 hpi) or IM *Cxcl10* (4 hpi) relative to uninfected macrophages.

Gene module activity based on new RNA levels for each single macrophage was quantified using Seurat’s *AddModuleScore()* function, and its association with *Salmonella* SPI2 activity was assessed by Spearman correlation with GFP fluorescence levels recorded during cell sorting.

#### Integrated Dynamo and CellRank2 analysis

To investigate gene expression dynamics across infection trajectories over the first 10 hours of infection, we computed RNA velocities based on new and old RNAs with Dynamo^23^ (v1.3.2). We used the standard preprocessing workflow for Seurat objects and “one-shot” type of RNA metaboling labeling experiments and ensured that all HVGs (from new, old and total RNA levels) selected with Seurat and DEGs from identified gene modules (Figure 3A) were retained.

Similarly, we formatted and pre-processed the Seurat object for CellRank2^24^ v(2.0.1). We used the RNA velocities computed with Dynamo to set up a Velocity kernel and the collection timepoints of each single macrophage to set up a Real Time kernel. We used a combination of both kernels as input for CellRank2 to compute the cell state transition matrix. We used a Generalized Perron Cluster Cluster Analysis (GPCCA) estimator to analyse the transition matrix, identify terminal states and compute cell fate probabilities. We selected all uninfected macrophages to define the initial state while IM *F10* and AIM *Adora3* were identified as putative terminal states by CellRank2. Finally, we leveraged diffusion pseudotime^52^ and the CellRank cell state transition matrix to construct a RNA velocity-informed pseudotime and order macrophages along trajectories towards Inflammatory and Anti-inflammatory states.

#### Gene module expression dynamics along trajectory pseudotime

We chose the uninfected macrophage population as our reference baseline to visualize gene expression dynamics. Gene module scores computed on new RNA levels for each macrophage (see “Gene module definition and correlation with *Salmonella* SPI2 activity” section) were centered by subtracting the mean gene module score calculated in mock (uninfected) macrophages.

Cells were ordered according to CellRank2 pseudotime and partitioned into 20 equal-sized bins (22 or 23 cells per bin). Weighted mean module scores were computed within each pseudotime bin using CellRank-derived fate probabilities as weights. Specifically, for each module and bin, weighted averages were calculated separately for each trajectory using the corresponding fate probability (IM *F10* or AIM *Adora3*) as weights.

Modules were grouped into pairs corresponding to early and late components within each module class (A–D), and their dynamics were jointly visualized along trajectory pseudotime. For the AIM Adora3 trajectory, the final pseudotime bin was excluded for visualization as it contained cells exclusively committed to IM F10. Smoothed trends were obtained using local regression (LOESS) to facilitate visualization of temporal patterns in module activity.

#### Differential regulation of gene modules and cell fate specification

We chose the IM *Cxcl10* (4hpi) macrophage state as our reference baseline to investigate gene module dynamics beyond the early response. Gene module scores computed on new RNA levels for each macrophage (see “Gene module definition and correlation with *Salmonella* SPI2 activity” section) were standardized relative to IM *Cxcl10* macrophages by calculating z-scores using the mean and standard deviation of each module within this reference state.

Cells from selected macrophage states in Figure 3C were retained for analysis. For Intermediate macrophages, CellRank-derived dominant fate assignments as defined in Figure 3B were used to further stratify cells according to their predicted trajectory.

For each group, average standardized module scores were computed and statistical significance was assessed using one-sample t-tests comparing z-scores to zero (corresponding to the reference IM *Cxcl10* state), followed by Benjamini–Hochberg correction for multiple testing. Adjusted p-values were used to annotate significance levels.

#### Upstream TF inference with CollecTri

Gene regulatory networks were reconstructed using the CollecTRI^25^ database of curated mouse transcription factor-target interactions. For each DEG from all four modules defined previously, upstream regulatory paths were recursively identified using directed graphs implemented in igraph (v2.0.3). The resulting subnetworks were simplified by removing transcription factors with fewer than four downstream interactions and target genes lacking upstream regulators. Separate inflammatory and anti-inflammatory regulatory networks were generated and subsequently merged into a combined network to identify shared and condition-specific regulators. Final network visualizations were generated with tidygraph (v1.3.0) and ggraph (v2.2.1) using Sugiyama hierarchical layouts.

### ATAC-seq/RNA-seq Data Analyis

#### Preprocessing of ATAC-seq data

Raw fastq files were generated by demultiplexing with bcl2fastq and general quality control was performed with FastQC v0.12.1 and MultiQC v1.15. Sequencing reads were trimmed with Cutadapt v4.4 to remove adapter sequences (CTGTCTCTTATACACATCT). Reads shorter than 20bp after trimming were excluded. Trimmed reads were aligned to the same combined reference genome as described above. Alignment was done with bowtie2 v2.3.4.1 (and the following additional parameters: -X 2000; –sensitive ; --no-mixed ; --no-discordant). Generated Bam files were processed to remove PCR duplicates with the MarkDuplicates function of Picard (v2.18.29), filtered for high-quality alignments with sambamba (v1.0.1) and mm39 blacklisted regions as well as reads mapping to mitochondrial chromosome or non-canonical chromosomes were filtered out with the intersect function of bedtools (v2.17.0) and samtools (v0.1.19), respectively.

Peak calling was done with MACS2 v2.2.9.1 (and the following additional parameters: -f BAMPE; -g mm ; --shift -37; --extsize 75 ; -p 1e-3 ; --nomodel ; --keep-dup all ; --call-summits). Reproducible peaks between the two biological replicates for each condition were identified with the Irreproducibility Discovery Rate (IDR) statistical framework (idr v2.0.4.2). Only peaks passing the IDR threshold of 0.05 were considered for further analysis and were annotated with the annotatePeaks.pl function from HOMER (v5.1) to connect each peak to the nearest gene in the reference genome.

#### Preprocessing of RNA-seq data

Raw fastq files were generated by demultiplexing with bcl2fastq and general quality control was performed with FastQC v0.12.1 and MultiQC v1.15. Sequencing reads were trimmed with Cutadapt v4.4 to remove the first 12bp of each read, poly-A tails and low quality bases (Phred score < 20). Reads shorter than 20bp after trimming were excluded. Trimmed reads were aligned to the same combined reference genome as described above. Alignment was done with STAR v020201. Reads mapping to exonic regions were quantified and summarized at the gene level for each library using *featureCounts* from Subread (v2.0.6). All *featureCounts* outputs were merged together to create the read count matrix.

#### Analysis of RNA-seq data

Genes with less than 100 read counts in half of the samples were considered lowly expressed and removed from the read count matrix. DEG testing was performed with DESeq2 (v1.30.1) using a model accounting for biological replicate batch effects. Genes with an adjusted p-value < 0.05 were considered differentially expressed (Wald test with Benjamini-Hochberg adjustment procedure).

For comparison with the scRNA-seq dataset, a read count matrix of pseudobulks was generated by aggregating read counts of single cells from macrophage states matching sampled conditions. Both matrices were merged together and processed with DESeq2 followed by principal component analysis for visualization after removal of batch effects associated with the type of data (bulk RNA-seq or pseudobulk) and biological replicates with the *removeBatchEffect* function from limma (v3.46.0).

#### Differential peak analysis and integration with RNA-seq

ATAC-seq peaks were called independently for each sample and analyzed using the DiffBind R package (v3.16.0). Differential chromatin accessibility between experimental conditions was assessed using the DESeq2 statistical framework implemented in DiffBind. Significantly differentially accessible regions were identified using a false discovery rate (FDR) threshold of 0.05. Differential peaks were annotated to nearby genes using ChIPseeker (v1.42.0).

To assess the relationship between changes in chromatin accessibility and gene expression, gene-associated peaks differentially accessible (ATAC-seq log fold change) and gene expression variations (RNA-seq log2 fold change) were compared. Combined significance was calculated as the product of adjusted p-values from both assays.

#### Transcription factor footprinting and TFBS count matrix generation

We used TOBIAS^30^ (v0.17.1) to perform TF footprinting analysis from the processed ATACseq dataset. Bam files from the two biological replicates were pooled together per condition. A merged peak file was created by combining all peaks retained after IDR analysis across all conditions. Tn5 transposase insertion bias was corrected with the ATACorrect function, footprinting scores were calculated with the ScoreBigwig function and differential binding analysis between and samples was done with the BINDetect function and JASPAR motifs^53^ (2024 core vertebrate motifs).

To generate the TFBS count matrix motif-specific overview files generated by the BINDetect function of TOBIAS were parsed for each experimental condition, and transcription factor binding sites were assigned to their corresponding ATAC-seq peaks. Binding sites simultaneously occupied in both SPI2-active WT and Δ*ssaV* at 18 hpi conditions were removed to focus on condition-specific binding events. For each transcription factor, the number of occupied binding sites per peak was calculated to generate TFBS-by-peak occupancy matrices.

#### TFBS count matrix analysis

TFBS occupancy count matrices were converted from peak-level to gene-level by summing occupancy scores across all regulatory peaks assigned to the same target gene. Analyses were restricted to genes significantly differentially expressed at the RNA level between SPI2-active WT and Δ*ssaV* at 18 hpi conditions and to TF motifs corresponding to transcription factors detected in bulk RNA-seq. Redundant motifs belonging to the same BINDetect motif cluster were merged by averaging their respective counts. After removal of low-information genes, matrices were log-transformed and scaled. Principal component analysis was performed independently for each of the four conditions (Uninfected, 2 hpi, Δ*ssaV* at 18 hpi, and SPI2-active WT at 18 hpi), and the top 6 principal components were integrated into a joint feature space. Gene clusters were identified with the Leiden algorithm^51^ on a nearest-neighbor graph (RANN, v2.6.2) and visualized by UMAP.

### scCRISPR / CROP-seq data analyis

#### Pre-processing of sequencing data

Raw sequencing data for all three types of libraries (GEX, MULTI-seq, CROP-seq) were demultiplexed and converted to FASTQ files with Cell Ranger (10x Genomics, v9.0.1) with the mkfastq command.

For the GEX libraries, sequencing reads were mapped to the mouse reference genome GRCm39 (mm39) to generate bam files and gene count matrices with the Cell Ranger count command. Bam files were converted to CIT files with the GEDI toolkit and single cells were assigned their 4sU labeling status based on MULTI-seq data. New RNA were quantified and NTR matrices were computed using GRAND-SLAM (v3.0.5)^19^ followed by conversion to Seurat Object format with grandR^54^ (v0.2.6).

FASTQs from MULTI-seq libraries were processed with deMULTIplex2 (v1.0.2)^55^ to extract MULTI-seq barcode counts associated with each cell barcode and assign each cell to its sample of origin.

FASTQs from CROP-seq libraries were processed with cutadapt to retain only read pairs with Read 2 containing the U6 promoter sequence (CTTGTGGAAAGGACGAAACACC). Read 2 was also trimmed to start with the captured sgRNA sequence. Trimmed reads were processed with deMULTIplex2 to extract sgRNA counts associated with each cell barcode and to perform sgRNA assignment per cell.

#### Dimensional reduction and clustering based on new and old RNAs

Analysis was conducted in R with the Seurat package (v4.0). Genes detected in fewer than 3 cells were filtered out and cells with more than 20,000 UMIs or less than 1,500 UMIs were excluded from analysis. Cells identified as multiplets based on MULTI-seq data were also removed. Cell cycle phase was inferred from log-normalized total RNA counts with Seurat’s CellCycleScoring function.

For joint analysis of new and old RNA, only 4sU-labeled cells were included. Both matrices were processed in parallel, including normalization and log-transformation of UMI counts, selection of the top 800 highly variable genes, and scaling with regression of biological replicate effects. Principal component analysis was performed using irlba (v2.3.5.1). Then, a joint PCA matrix was built by combining the top 15 PCs from each assay (new and old) before dimensional reduction with UMAP. The joint PCA was also used to build a KNN graph (k = 20) followed by cell clustering with the Leiden algorithm^51^. We used the C implementation for R of the Leiden algorithm (leidenbase v0.1.3, Trapnell Lab) with a resolution set at 0.002.

#### sgRNA efficiency assessment and perturbation enrichment analysis

To evaluate sgRNA knockdown efficiency, cells were grouped by sgRNA and their transcriptomes were aggregated and normalized to counts per million (CPM). Target gene expression levels were compared to those in cells expressing non-targeting control (NTC) sgRNAs to compute fold changes. sgRNAs that reduced target gene expression by at least 40% relative to NTCs were classified as effective. Guides with low representation across the dataset (< 5 cells) were excluded from analysis.

Enrichment of specific gene knockdowns relative to NTCs in inflammatory or anti-inflammatory macrophage states was evaluated by logistic regression using a generalized linear model (GLM) with a binomial distribution. To ensure robust estimation, only gene knockdowns represented by at least 10 perturbed cells (i.e. cells harboring effective sgRNAs) were included. Target gene and biological replicate were included in the model as explanatory variable and covariate, respectively. Model coefficients were converted to odds ratios, and statistical significance was assessed using Wald tests. P-values were adjusted for multiple testing using the Benjamini–Hochberg procedure. Target genes with FDR ≤ 0.1 were considered as significantly enriched.

#### Differential gene expression analysis of CRISPR perturbations

Differential gene expression upon CRISPR perturbation was assessed using SCEPTRE^36^ (v0.10.3). Analyses were performed both across all macrophage states (naive, inflammatory, and anti-inflammatory) and separately within each state to identify shared and condition-specific perturbation effects. Only gene knockdowns represented by at least 10 perturbed cells per condition were included.

Analyses were conducted using the total RNA count matrix after filtering out lowly expressed genes (detectable expression in less than 5% of cells). The number of captured UMIs, number of detected genes, percentage of mitochondrial UMIs, biological replicate, cell cycle phase, macrophage state (when necessary) and global percentage of new RNA were included as covariates. The recommended SCEPTRE workflow was followed for low MOI experiments, cell quality control, calibration and power checks and running the discovery analysis.

Given the limited number of cells per perturbation and the resulting constraints on statistical power, a relatively permissive false discovery rate (FDR ≤ 0.2) was used to prioritize sensitivity. Identified hits were interpreted in the context of effect size and consistency across conditions.

To determine if SCEPTRE hits were also supported at the new RNA level, we used the log-transformed and normalized new RNA count matrix to compute effect sizes as the difference in mean expression between perturbed and NTC cells. Statistical significance was assessed using a Wilcoxon rank-sum test, and p-values were adjusted for multiple testing using the Benjamini–Hochberg procedure. Interactions with FDR ≤ 0.1 were considered supported by nascent RNA levels.

## ACKNOWLEDGMENTS

The authors thank Nina DiFabion (Saliba lab) for support in sequencing, Redmond Smyth and Jianhui Li (IBMC, Strasbourg, France) for Nanopore sequencing of sgRNA library, Thomas Rudel (Biocenter, University of Wuerzburg, Germany) for kindly providing Hoxb8 progenitors originally created by Dr. David Sykes (Massachusetts General Hospital, Boston, USA), Florian Erhard (University of Regensburg) for support on scSLAM-seq analysis, and the Beisel lab (HIRI) for molecular work support.

We thank the funding received from the Bundesministerium für Forschung, Technologie und Raumfahrt CompLS – Computational Life Sciences HOPARL (031L0289B) for A.-E.S., C.T. and F.T. and from the Deutsche Forschungsgemeinschaft (DFG, German Research Foundation) – Cluster for Nucleic Acid Sciences and Technologies – NUCLEATE (Project-ID 533767322 – EXC 3113/1) to A.-E.S, F.T. and C.T. and DFG-funded SFB 1583 Decisions in Infectious Diseases (DECIDE; Projektnummer #492620490 B05 and Z02) to A.-E.S and C.T. C.T. acknowledges kickoff funding through the HIRI graduate training program “RNA & Infection.” B.K. thanks the Munich School of Data Sciences (MUDS) for Ph.D. fellowship support. P.W.S.H is supported by a Wellcome Trust Career Development Award (226546/Z/22/Z).

---

**Figure S1:**
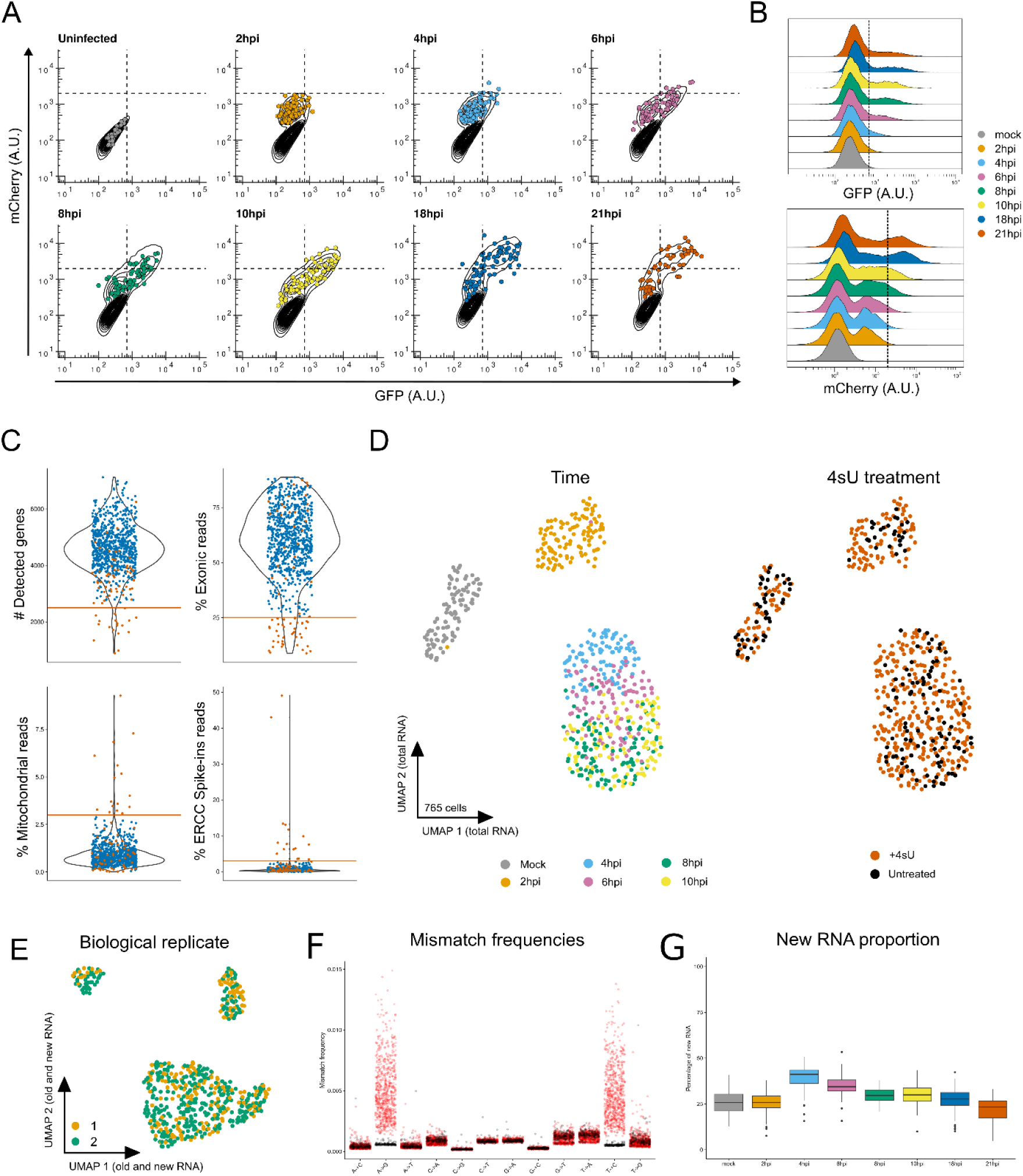
Salmonella SPI2 induction dynamics and general quality controls of the scSLAM-seq dataset, related to Figure 1. **(A)** Flow cytometry scatter plots displaying intracellular Salmonella SPI2 activity of uninfected and infected macrophages, split by infection time points. Contour lines correspond to flow cytometry data recorded at the population level, and colored dots indicate each single macrophage collected for downstream analysis with scSLAM-seq. **(B)** Ridgeline plots showing the intracellular Salmonella fluorescence intensity distribution of mCherry (top) and GFP fluorescence (bottom) across time points of infection. Dashed lines correspond to thresholds used to determine Salmonella growth status and SPI2 activity (Figure 1B **and Suppl.** Fig. 1A). **(C)** Violin plots showing the general quality metrics for all single cells analyzed by scSLAM-seq and filtering criteria for sorting out low-quality cells. Orange solid lines represent the selected threshold for the indicated criteria. Single cells filtered out based on any criteria are represented in red. **(D)** UMAP visualization of 765 single-macrophage transcriptomes obtained with scSLAM-seq, including 187 cells not exposed to 4-thiouridine (4sU). UMAP visualization results from an analysis solely based on total RNA levels. Single macrophages are colored by timepoint of collection (left) or 4sU treatment status (right). **(E)** UMAP visualization of 578 single-macrophage transcriptomes obtained by scSLAM-seq as in Figures 1C and **1D** and colored-coded by biological replicate. **(F)** Nucleotide mismatch frequencies calculated with GRAND-SLAM in 4sU-treated single cells (red) and untreated cells (black). T-to-C mismatches correspond to sense reads i.e. the original mRNA sequence, while A-to-G mismatches correspond to antisense reads, i.e. the reverse-complement of mRNA sequence. **(G)** Distribution of global new RNA percentages per cell as computed by GRAND-SLAM and stratified by time point of collection. Box plots denote the median (centre line) and interquartile range (box), and whiskers extend to the most extreme values within 1.5× of the interquartile range. Observations outside this range are plotted as individual outliers.

**Figure S2:**
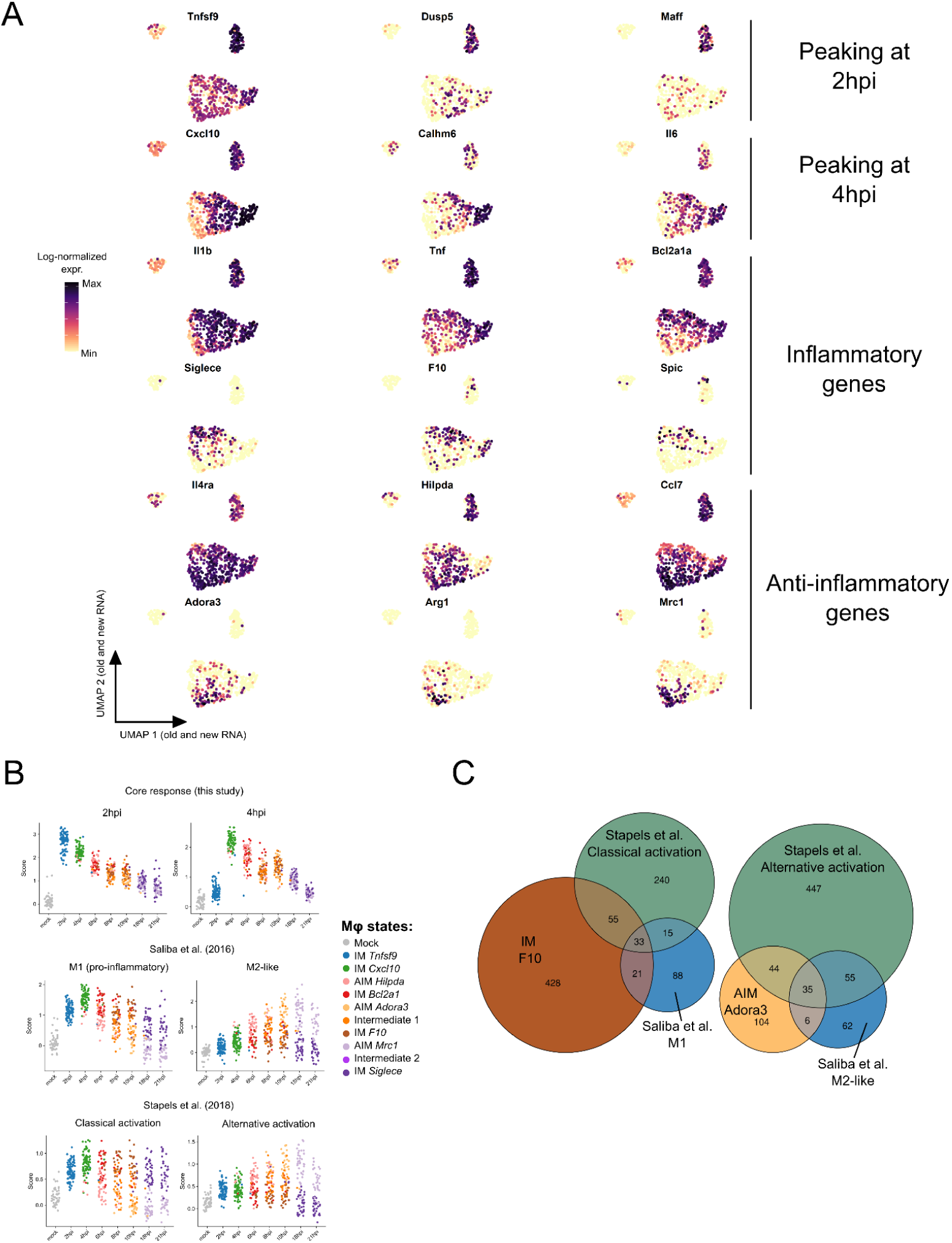
Inflammatory and anti-inflammatory macrophage polarization signatures, related to Figure 2. **(A)** Log-normalized read counts from new RNA levels of selected marker genes, color-coded and projected onto the UMAP embedding in Figure 2A. **(B)** Gene set module scores of early response (DEGs from 2 hpi and 4 hpi) from Figure 2C and M1/M2 macrophage signatures derived from two published datasets (Saliba et al., 2016; Stapels et al., 2018). Scores were calculated based on new RNA levels for each of the 578 single macrophages. Data points are grouped per timepoint of collection and colored by macrophage state as indicated in Figure 2A.

**Figure S3:**
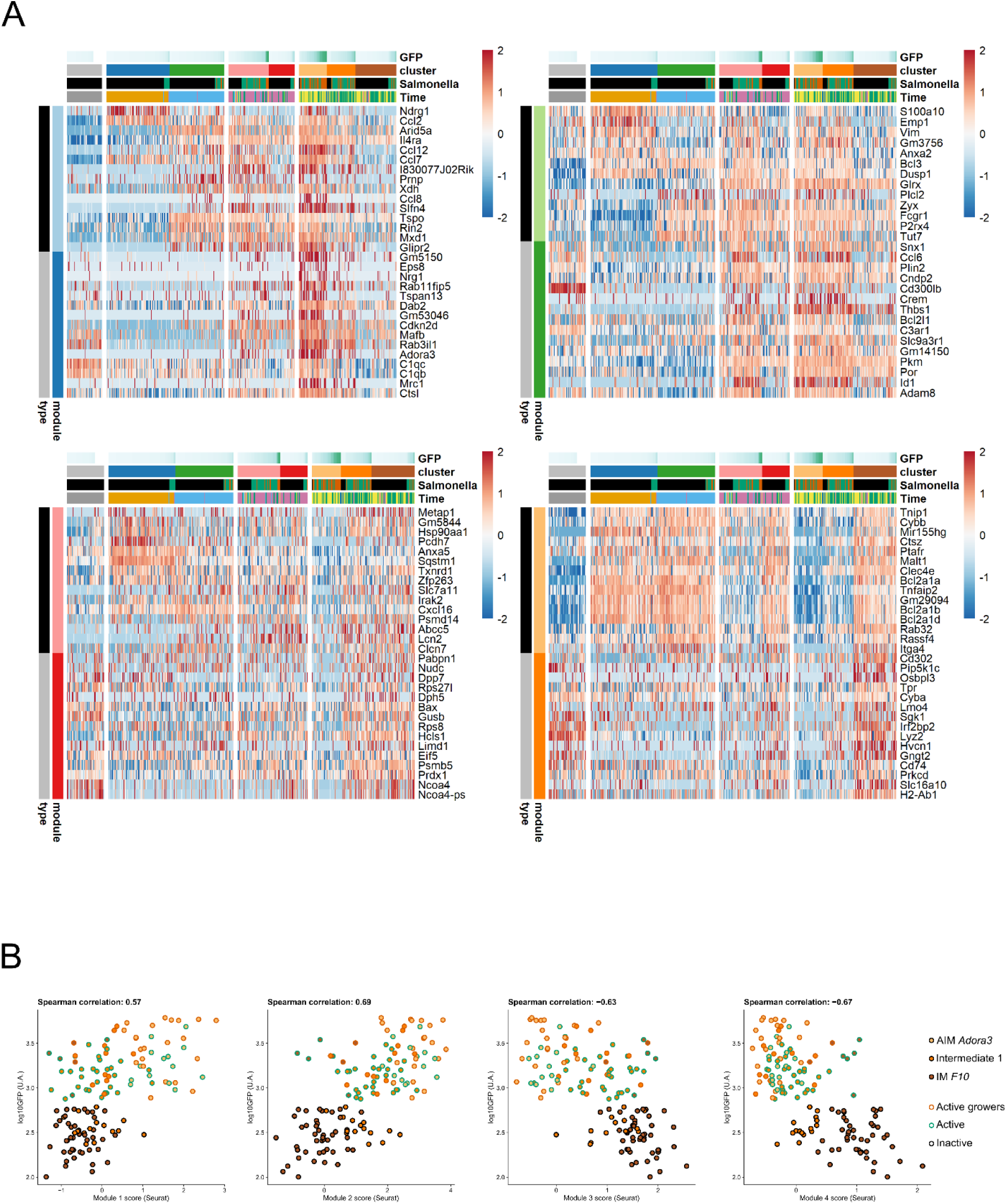
Differential expression of gene modules linked to Inflammatory and Anti-inflammatory signatures is correlated to Salmonella SPI2 activity, related to Figure 3. **(A)** Heatmaps displaying z-scores of log-normalized new RNA readcounts of selected differentially expressed (DE, FDR < 0.05) genes from each module defined in Figure 3A (top 15 genes with lowest p-values). Cells within each macrophage state, i.e. columns, are ordered based on measured GFP fluorescence levels (increasing GFP signal from left to right). Statistical significance of differential expression for each gene per macrophage state is shown in Table S2. **(B)** Spearman correlation between main gene module scores computed on log-normalized new RNA readcounts and GFP fluorescence levels measured during single-cell sorting. The analysis is restricted to Inflammatory, Anti-inflammatory and partially reprogrammed macrophage states (n = 135 cells). Data points are colored by macrophage state as indicated in Figure 2A and border colors indicate Salmonella status as defined in Figure 1B.

**Figure S4:**
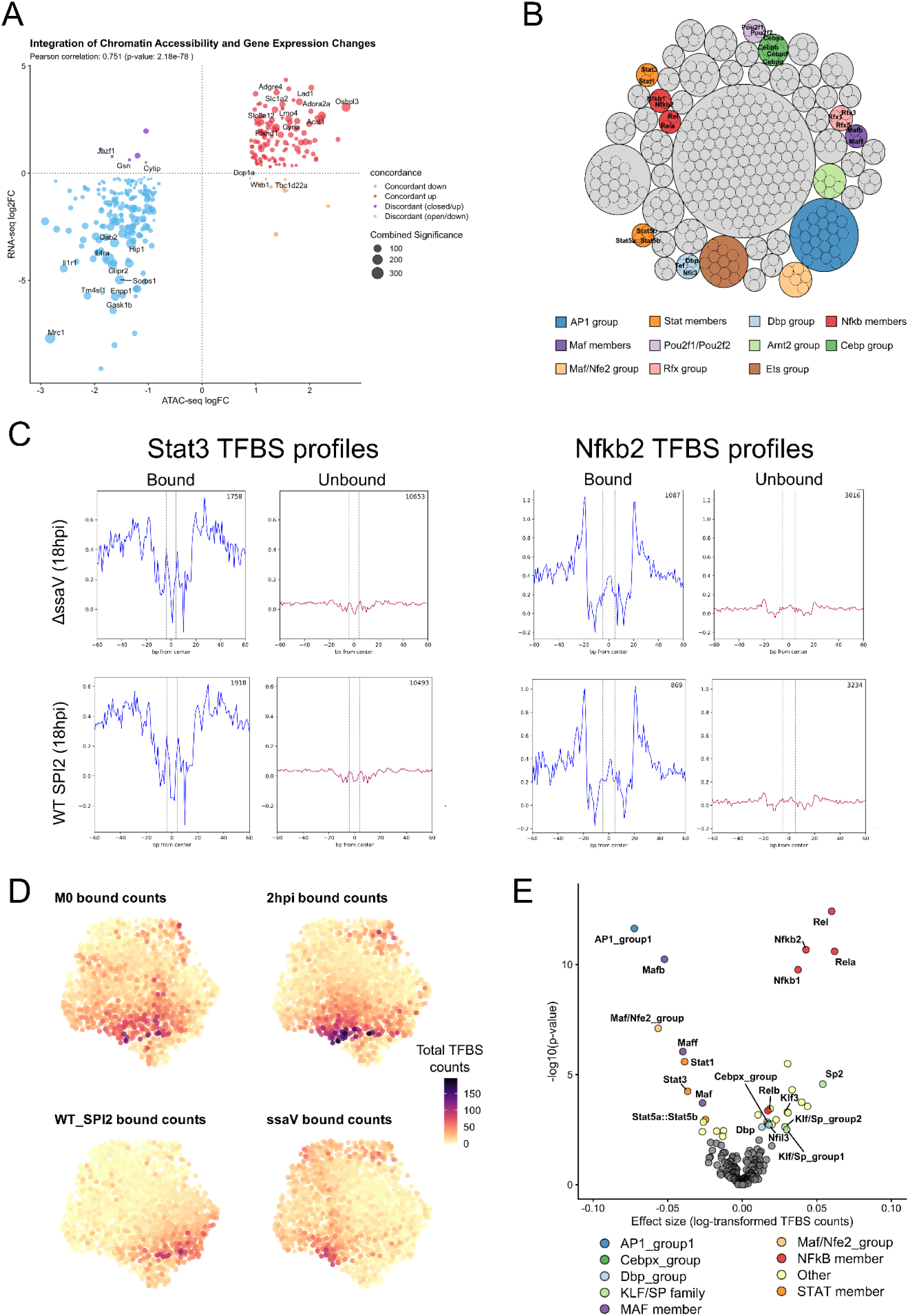
Complementary analyses of TFBS occupancy at the gene level across conditions enhances TF activity profiling, related to Figure 5. **(A)** Scatter plot showing the relationship between differential chromatin accessibility (ATACseq) and gene expression (RNAseq). Each point represents one gene, plotted by log2 fold change (log2FC) in chromatin accessibility (ATAC-seq, analysis done with DiffBind) versus log2FC in gene expression (RNA-seq, analysis done with DESeq2), both computed for macrophages infected by ΔssaV mutant versus SPI2-active WT Salmonella. Only genes differentially expressed (FDR < 0.05, Wald test adjusted with Benjamini–Hochberg correction) and differentially accessible were included (FDR < 0.05, Wald test adjusted with Benjamini–Hochberg correction). Point color denotes concordance between chromatin accessibility and gene expression changes, point size reflects combined statistical significance (−log10 of the product of adjusted p-values), and labeled genes correspond to top-ranked features based on an integrated score combining effect size and significance across both datasets. **(B)** Circle packing representation of JASPAR motifs grouped by TOBIAS clustering. Motifs are clustered based on the overlap of predicted TF binding sites across all peaks. Only motifs associated to TFs expressed at the RNA level were retained. Selected motifs of interest are labeled. Colors reflect TF groups highlighted in Figure 5B. **(C)** Aggregated footprint plots for Stat3 and Nfkb2 centered on predicted binding sites (dashed lines). For each TF, the footprint plots are splitted by TF bounding status and experimental condition. Total numbers of TF binding sites considered for each plot are indicated (top-right corners). **(D)** Raw bound TFBS counts per gene across four conditions, color-coded and projected onto the UMAP embedding in Figure 5C. **(E)** Pairwise comparison of TF binding site (TFBS) enrichment between gene clusters preferentially associated with the ΔssaV mutant or SPI2-active WT Salmonella conditions. The volcano plot displays the difference in mean log-transformed TFBS counts between DEGs upregulated in ΔssaV-infected BMDMs and DEGs upregulated in SPI2-active WT–infected macrophages against the −log10(p-value) derived from a Wilcoxon rank-sum test. Each point represents a single JASPAR motif or group of JASPAR motifs (based on overlapping motif sites). TFBS counts for each cluster were computed from the condition in which that cluster shows maximal TFBS occupancy, enabling comparison of condition-specific binding signatures between gene groups. Motifs significantly enriched are highlighted (FDR < 0.05, Wilcoxon rank-sum test with Benjamini–Hochberg correction) and colored by family.

**Figure S5:**
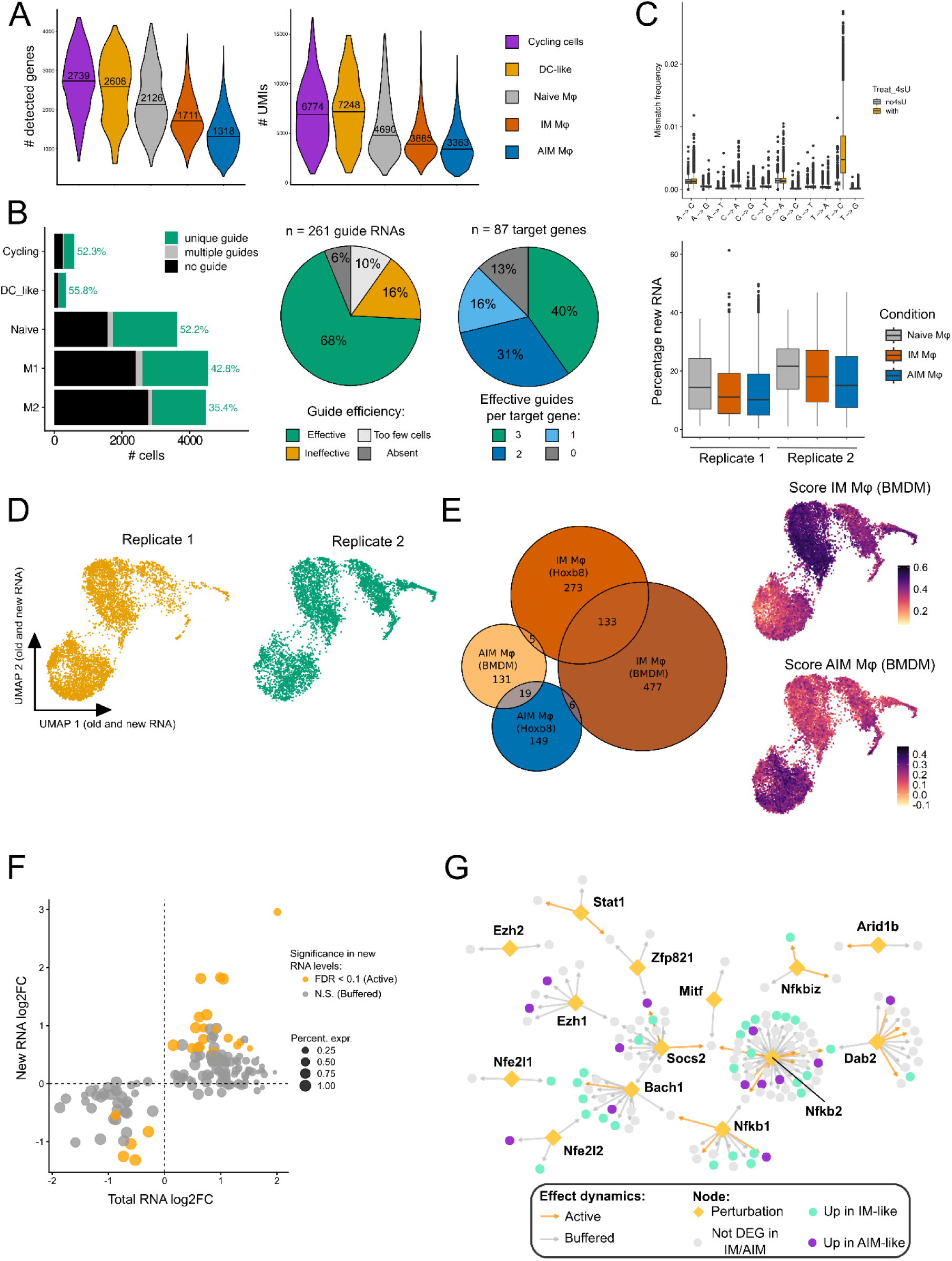
Quality control, perturbation efficiency, and RNA dynamics analyses in Hoxb8 macrophages, related to Figure 6. **(A)** Gene expression metrics in Hoxb8 cells passing quality control. Violin plots for number of detected genes per single cell (left) and total numbers of UMIs captured (right). Violins are filled with color matching cell states as in Figure 6B. Displayed numbers and lines in violins correspond to the median per cell state. **(B)** Single-cell CRISPR metrics. (Left) Assignment of guide RNA identity for each of the 11,758 Hoxb8 macrophage transcriptomes. Each bar corresponds to a cell state as in Figure 6B. Bar heights indicate the total number of cells. Bar colors indicate guide assignment status. Displayed percentages correspond to the proportion of single-cells associated to a unique guide RNA per cell state. (Middle) Pie chart indicative of each guide RNA efficiency to repress their target gene. Guide RNAs resulting in at least 40% of target gene expression relative to NTCs were considered effective. Efficiency was not assessed for guide RNAs observed less than 5 times across the whole dataset for a given guide. (Right) Pie chart summarizing the number of effective guide RNAs per targeted gene. **(C)** RNA metabolic labeling metrics. (Top) Boxplots of nucleotide mismatch frequencies obtained with GRAND-SLAM 3.0 in 4sU-treated single cells (orange) and untreated cells (grey). (Bottom) Boxplots of global new RNA percentage per cell as computed by GRAND-SLAM 3.0. Cells are grouped per sample identity and biological replicate. **(D)** UMAP embedding of 11,758 Hoxb8 macrophage transcriptomes obtained by scSLAMseq 2.0. UMAP embedding results from the joint analysis of new and old RNA levels. Single macrophages are colored by biological replicate. **(E)** Overlap between Hoxb8 macrophages and BMDM polarization signatures. (Left) Venn plot of differentially expressed genes (FDR < 0.05, Wilcoxon Rank Sum test with Benjamini-Hochberg correction) marking Inflammatory and Anti-inflammatory signatures in BMDM (Figure 2C) and Hoxb8 macrophages. Circle size is proportional to the total numbers of DEG within each group. Indicated numbers correspond to the number of specific or shared DEGs between groups. (Right) Signature module scores of Inflammatory and Anti-inflammatory BMDMs (Figure 2C) projected on the UMAP embedding. Module scores were calculated based on total RNA levels. **(F)** Concordance plot of perturbation effects between total and new RNA levels, as reported in Figure 6D. Each point represents a significant perturbation effect, defined as a gene targeted by a guide RNA and a downstream gene affected by the perturbation. The x-and y-axes indicate the log2 fold change in downstream gene expression relative to non-targeting control (NTC) cells for total and new RNA levels, respectively. Point size reflects the percentage of cells with detectable expression of the downstream gene in total RNA. Perturbation effects that are also significant at the new RNA level are highlighted (FDR < 0.1, Wilcoxon test with Benjamini–Hochberg correction). **(G)** Gene perturbation network as in Figure 6E with edge colors indicating whether the perturbation effect is detected only in total RNA levels (Buffered) or in both total and new RNA levels (Active).

